# Ultra-sensitivity metaproteomics redefines the gut “dark metaproteome”, uncovering host-microbiome interactions and drug targets in intestinal inflammatory diseases

**DOI:** 10.1101/2024.04.22.590295

**Authors:** Feng Xian, Malena Brenek, Christoph Krisp, Doriane Aguanno, Elisabeth Urbauer, Tharan Srikumar, Ranjith Kumar Ravi Kumar, Qixin Liu, Allison M Barry, Bin Ma, Jonathan Krieger, Dirk Haller, Manuela Schmidt, David Gómez-Varela

## Abstract

The gut microbiome is a complex ecosystem with significant inter-individual variability determined by hundreds of low-abundant species as revealed by genomic methods. Functional redundancy demands direct quantification of microbial biological functions to understand their influence on host physiology. This functional landscape remains unexplored due to limited sensitivity in metaproteomics methods. We present uMetaP, an ultra-sensitive metaproteomic solution combining advanced LC-MS technologies with a novel FDR-controlled de novo strategy. uMetaP improves the taxonomic detection limit of the gut "dark metaproteome" by 5,000-fold with exceptional quantification precision and accuracy. In a mouse model of colonic injury, uMetaP extended metagenomics findings and identified host functions and microbial metabolic networks linked to disease. We obtained orthogonal validation using transcriptomic data from biopsies of 204 Crohn’s patients and presented the concept of a "druggable metaproteome". Among the drug-protein interactions discovered are treatments for intestinal inflammatory diseases, showcasing uMetaP’s potential for disease diagnostics and data-driven drug repurposing strategies.

## INTRODUCTION

The gut microbiome, a complex ecosystem of hundreds of bacterial species, plays a crucial role in host physiology, affecting overall health^1^. While a core set of microbial species is shared among most individuals, significant variability exists due to medium- and low-abundance taxa^2, 3, 4^. This variability contributes to personalized microbiomes and challenges the concept of a unique healthy microbiome^5^. Although genomic methods have greatly expanded our understanding of the taxonomic repertoire, functional redundancy among microbiome members requires methods that can directly quantify the biological functions of the microbiota and host.

Metaproteomics, which analyzes microbial samples using liquid chromatography coupled with mass spectrometry (LC-MS)-based proteomics, has emerged as a powerful tool for investigating the functional signatures of host-microbiome interactions in health and disease^6^. However, over 80% of bacterial species detected by genomic methods remain undetected by metaproteomics, constituting the "dark metaproteome"^4^. Significant improvements in the sensitivity of metaproteomic approaches are needed to explore the highly complex and largely uncharted functional landscape of the gut microbiome. We present uMetaP, an integrative, ultra-sensitive metaproteomic solution that achieves exceptional depth and sensitivity in studying complex metaproteomes.

uMetaP combines Ultra-High-Performance Liquid Chromatography (UHPLC), an optimized ionization source to maximize ion transfer^7^, and the sensitivity of the timsTOF Ultra mass spectrometer^8, 9^. Using mouse feces as a model, uMetaP fragmented over 1.6 million precursors via Data-Dependent Acquisition Parallel Accumulation–Serial Fragmentation (DDA-PASEF). However, less than 30% resulted in confident peptide spectrum matches (PSMs). We trained a de novo algorithm, Novor^10^, on 1.7 million PSMs, marking the first instance of a de novo algorithm trained in PASEF’s four-dimensional data structure. We combined it with a multi-tier filtering procedure to enhance peptide confidence, enabling us to develop novoMP: a de novo-assisted metaproteomic database construction method. NovoMP expanded a mouse fecal metaproteomic database from 223 to 774 microbial species, including archaea, fungi, and viruses. The final database, with 208,254 microbial protein sequences (a 19-times increase from our previous PASEF-based database^11^), is available via PRIDE for community use.

When powered by Data-Independent Acquisition (DIA-) PASEF, uMetaP identified and quantified 210,051 microbial peptides and 118,937 microbial protein groups, tripling the previous state-of-the-art^11^. An orthogonal FDR control strategy ensured de novo-derived peptides matched traditional database identification confidence. uMetaP identified 1,043 proteins of unknown function^12^ (PUFs), 2,342 small proteins^13, 14^, and 581 antimicrobial peptides^15^ (AMPs). Using SILAC-labeled bacteria, we determined the accurate limit of detection and quantification for the gut "dark metaproteome", down to 0.0003% and 0.0044%, respectively, improving previous standards^4^ by 5,000-fold and enabling identification of previously undetectable low-abundance taxa.

uMetaP extended taxonomic changes observed by metagenomics on a transgenic mouse model of colonic injury. Further, we identified 990 host-regulated proteins and 92 microbiota-specific networks, revealing novel pathways in tissue damage. Orthogonal validation with Crohn’s patient transcriptomic data confirmed the regulation of 490 proteins. Using additional mouse transcriptomic data, 33 proteins showed consistent alterations across datasets linked to inflammation, metabolic functions, and mitochondrial activity. Network analysis highlighted protein hubs influencing tissue injury. We introduced the concept of the "druggable metaproteome", identifying 204 drug-protein interactions, including current therapies for inflammatory diseases, and offering resources for drug repurposing.

By integrating the latest LC-MS technology and a new de novo analysis strategy, as well as a transgenic mouse model of colonic injury, orthogonal validation using patient’s transcriptomic data, and a detailed drug-gene analysis, we show the potential of uMetaP in microbiome research. This includes uncovering functional signatures of health and disease and guiding new therapeutic interventions.

## RESULTS

### uMetap enables novoMP: a novel de novo sequencing strategy improving metaproteomic database construction

Our previous work introduced the benefits of Parallel Accumulation Serial Fragmentation (PASEF) in metaproteomics, including during the construction of a metaproteomic database^11^. Remarkably, analysis of the same eight peptide fractions using the new technological solutions integrated into uMetaP enabled the fragmentation of 4 times more precursor ions than when using our previous workflow based on a timsTOF Pro mass spectrometer (Figure 1A), resulting in 4 times more identified peptides (129,425 vs. 30,460; Supp. Figure 1A) and a significant shift towards higher peptide intensities (Supp. Figure 1B). Despite this considerable improvement, the classical database search identified fewer than 30% of the precursors fragmented in the timsTOF Ultra mass spectrometer (Figure 1A), leaving most biological data uncharacterized. We hypothesised that a de novo search strategy, which does not rely on a target sequence database, could rescue part of this valuable information. However, to our knowledge, no published de novo algorithms are trained in the 4-dimensional data structure of PASEF. Moreover, previous studies applying de novo for metaproteomic database construction lacked methodologies to test the confidence of peptide assignments^16^. This is especially critical in metaproteomics due to the immense peptide landscape of these complex samples^17^. We constructed novoMP, a novel strategy integrating the first algorithm, to the best of our knowledge, trained in PASEF data structure, together with a multi-layered quality control filtering strategy to rigorously select high-confidence de novo peptide-spectrum matches (PSMs; see Methods for details).

**Figure 1:**
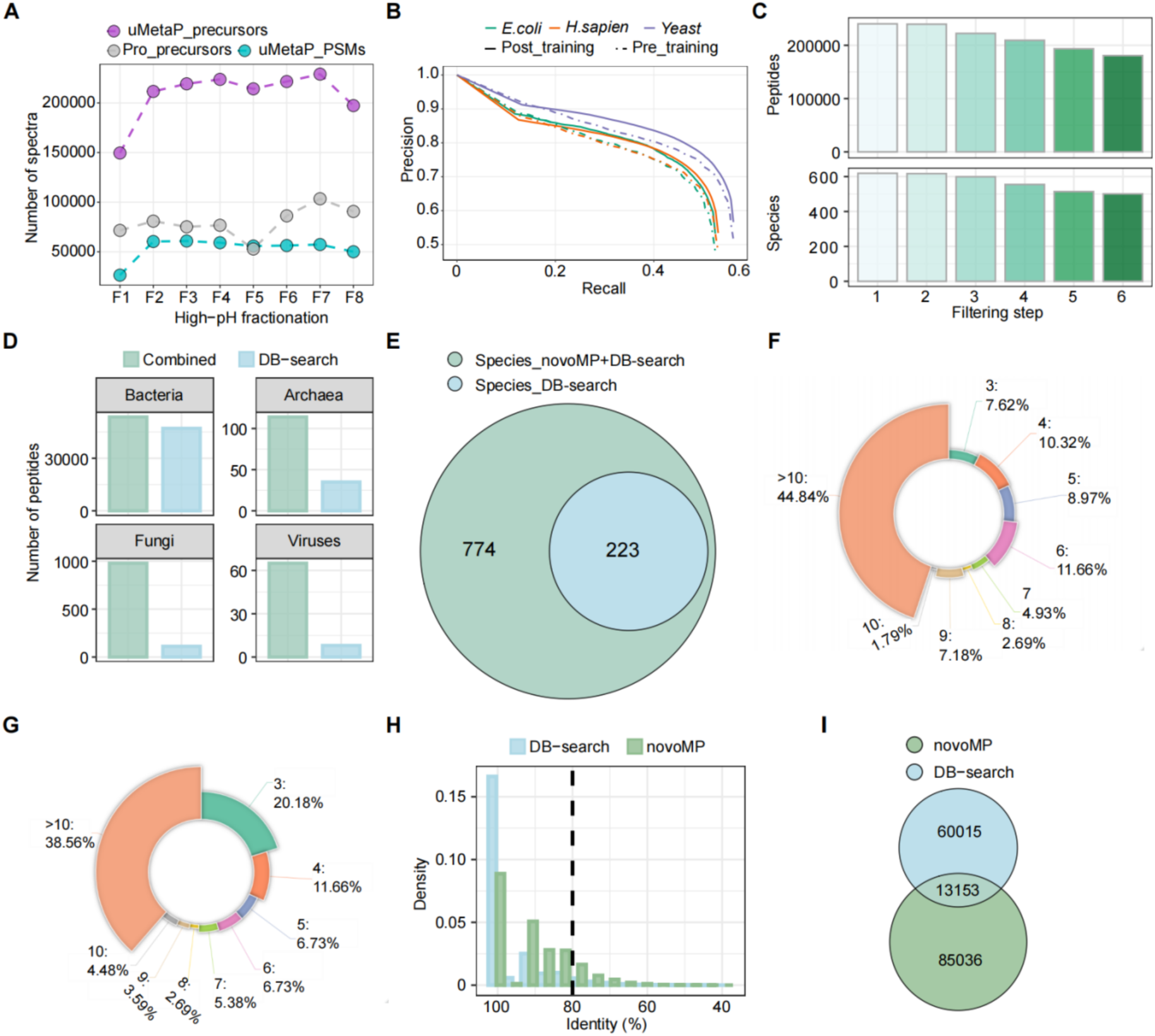
Development and impact of novoMP on metaproteomic database construction. (A) Comparison of the number of precursor ions fragmented and the resulting identified peptide- spectrum matches between the uMetaP workflow and our previous workflow across eight high-pH fractions. (B) Precision-recall curves illustrating the performance of the Novor algorithm on *E. coli*, *H. sapiens*, and Yeast datasets before (pre-) and after (post-) training with PASEF datasets. (C) Reduction in the number of peptides and species identified as filtering steps progress in the novoMP workflow. (D) Taxonomic coverage comparison of peptides annotated via the combined peptides (DB-search + novoMP) and DB-search peptides alone across bacteria, archaea, fungi, and viruses. (E) Venn diagram showing the unique and shared species identified using the novoMP integrated strategy versus DB-search alone. (F-G) Distribution of species-specific peptide counts for 223 species shared between Combined strategies (F) and DB-search (G). (H) Density plot of sequence identity percentages for BLAST+ homology searches against the NCBI RefSeq database using DB-search peptides and novoMP- derived peptides. A threshold of 80% (black-dotted line) sequence identity was used to filter high-confidence protein matches. (I) Venn diagram comparing protein sequences identified by DB-search and novoMP-derived peptides for the database construction.

We trained Novor^18^ using over 1.750,000 PSMs from PASEF data acquired on various timsTOF platforms (see Methods for details). The evaluation in a human-*E.coli*-yeast dataset not used during model training shows how the post-trained model maintains higher precision as recall increases compared to the pre-training model (Figure 1B; Supp. Figure 1C). These improvements result in an average of 5-7% gains concerning correct amino acid and peptide assignments in human, *E.coli*, and yeast peptides (Supp. Figure 1D). Similar improvements were found when samples were prepared with various enzymes (Supp. Figure 1E). Next, we applied this new de novo model to analyse pH-fractionated mouse fecal peptides acquired in Data-Dependent Acquisition (DDA)-PASEF. As a result of the multi-layered filtering strategy, unique novoMP peptides and annotated species counts decreased as the filtering steps progressed (Figure 1C and Supp. Figure 1F-1K). In comparison to taxonomy annotation using only peptides from classic database searches (DB-search), the integration of de novo peptides (Combined) improved taxonomic coverage, particularly for archaea, fungi, and viruses (Figure 1D). Of a total of 774 annotated species (Supp. Table 1) from all peptides (DB-search + uMetaP), only 223 species could have been identified by using solely DB-search peptides (aka. DB-search alone would have discovered a minimum of three species-specific peptides). Detailed analysis revealed the gains in taxonomic coverage reached by novoMP. For example, there is a marked increase in the number of peptides representing the above-mentioned 223 species when including de novo data (Figure 1F), compared to using DB-search peptides alone (Figure 1G). Moreover, the combination of peptides from DB-search + novoMP (Combined strategy) enabled the annotation of 551 additional species, increasing taxonomic coverage 247% (Figure 1E). Applying novoMP to archived DDA-PASEF data from our previous study^11^, increased the taxonomic coverage by 139 % (from 89 to 213 species; Supp. Figure 1L). The bigger gains enabled by novoMP in our new dataset, together with the remarkable taxonomic overlap among these independent sets of samples (Supp. Figure 1M; uMetaP discovers 90% of species from our previous study using a timsTOF Pro), demonstrated the benefit of novoMP to access valuable but otherwise hidden precursor information produced by the latest mass spectrometry technology.

Unlike DB-search, de novo sequencing does not inherently assign proteins to detected peptides. Thus, we conducted BLAST+ homology searches against the NCBI RefSeq database, applying the same pipeline to both novoMP-derived and DB-search-derived peptides. We set an 80% sequence identity threshold between query sequences and references to exclude low-confidence matches (Figure 1H). This approach retrieved 73,168 and 98,189 protein sequences for DB-search and novoMP-derived peptides, respectively, with 13,153 shared between the two (Figure 1I), totaling 158,204 unique protein sequences. Finally, we add 53,502 proteins identified through the classic DB-search against the mouse gut MGnify catalogue. As a result, we created a carefully curated mouse fecal metaproteomic database comprising 208,254 microbial protein sequences, which is available to the metaproteomic community via PRIDE.

In summary, the uMetaP workflow presented here demonstrated the power of the latest mass spectrometry instrumentation paired with a purpose-built de novo strategy for database construction, a critical step in metaproteomics.

### uMetaP powered by DIA-PASEF enhances taxonomic and functional coverage, sensitivity and quantitative precision

Our previous study introduced the benefits of combining Data-Independent Acquisition (DIA)- PASEF with deep neural network-based data analysis for complex metaproteomic samples^11^. uMetaP powered by DIA-PASEF increased 3 to 4 times the identifications of microbial and host peptides and proteins compared to our previous workflow^11^ (Figure 2A) when comparing similar conditions. Peptide identifications raised linearly and gradually plateauing at a total of 96,513 peptides (89,128 microbial and 7,385 mouse peptides; averaged across three replicates) when 25 ng of peptides were injected over a 30-minute gradient. Extending the LC gradient to 66 minutes further boosted the number of identified peptides and protein groups to 141,811 and 79,693, respectively (averaged across three replicates with 100ng peptide). Reflecting improved sensitivity, uMetaP detected an average of 200 microbial and 76 host protein groups at an ultra-low sample amount of 10 pg (Figure 2A and Supp. Table 2). uMetaP identified peptides spanning over four orders of magnitude using 25 ng of injected peptides with a 30-min gradient (Supp. Figure 2A) and showed a remarkable quantitative precision with more than 84% of peptides exhibiting a coefficient of variation (CV) lower than 0.2 (Supp. Figure 2A). In total, 210,051 microbial peptides were identified, with 32,400 of these added by novoMP to the mouse fecal metaproteomic database (novoMP-DB; Figure 2B).

**Figure 2:**
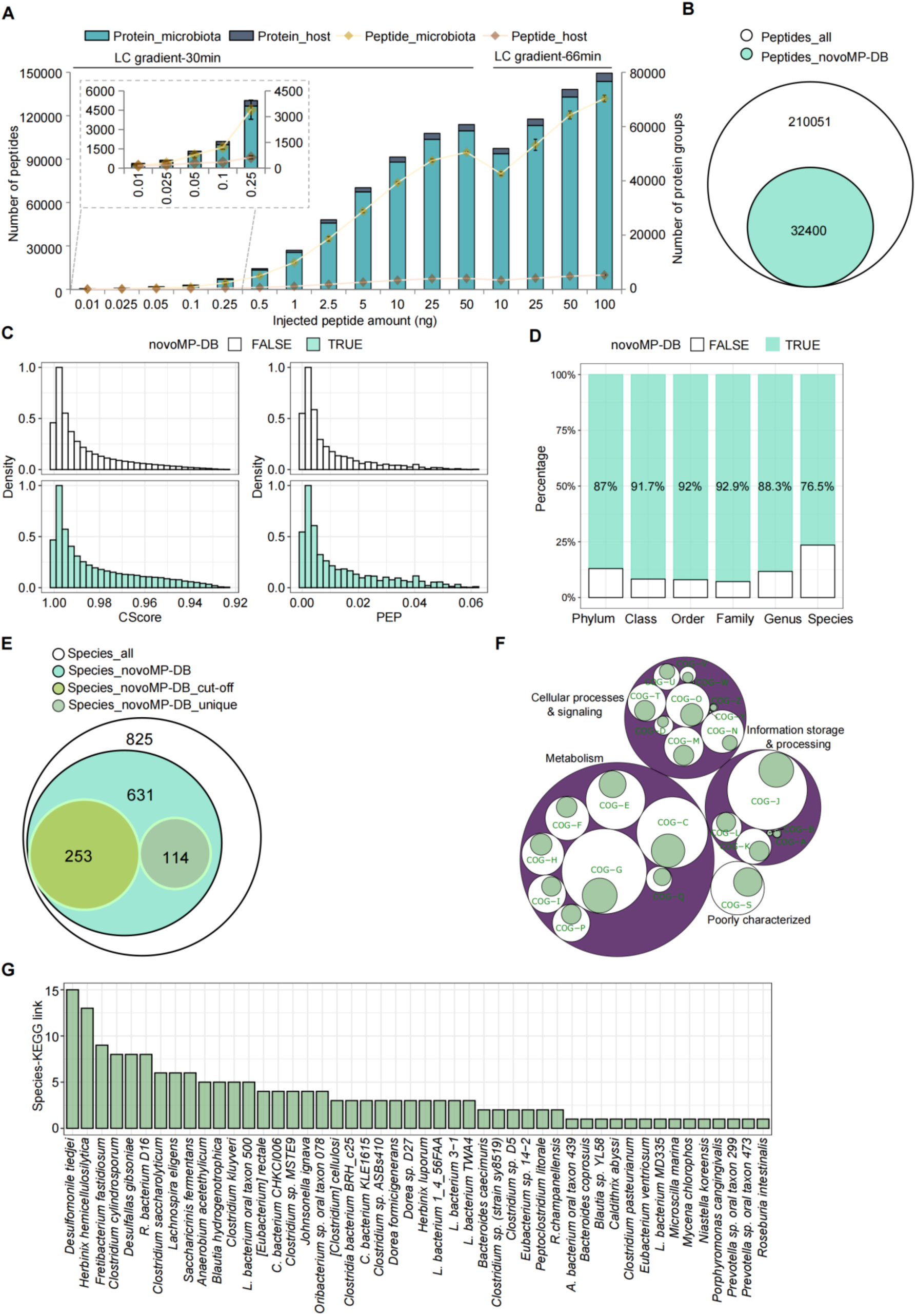
uMetaP powered by DIA-PASEF enhances metaproteome taxonomic and functional coverage, sensitivity, and quantitative precision. (A) Number of identified microbial and host peptides and protein groups using uMetaP powered by DIA-PASEF across varying sample amounts and LC gradient lengths. (B) Venn diagram showing the total microbial peptides identified, including 32,400 additional peptides added by novoMP to the metaproteomic database (novoMP-DB). (C) Distribution of CScore and Posterior Error Probability (PEP) values for peptides originated from novoMP-DB (“TRUE") and from DB- search (“FALSE”). (D) Taxonomic contributions of novoMP-DB peptides across ranks (phylum to species). (E) Venn diagram of species annotated using any type of peptide (Species_all), species with novoMP-DB peptides assigned (Species_novoMP), annotation meeting the 3- species-specific peptide cut-off due to the addition of novoMP-DB (Species_novoMP-DB_cut- off), and species uniquely annotated with novoMP-DB peptides (Species_novoMP- DB_unique). (F) Functional representation of protein groups mapped to Clusters of Orthologous Genes (COGs). The percentages of proteins uniquely discovered by novoMP-DB were shown in green. The white bubble area represents the percentages of functional clusters discovered by all detected proteins. (G) Species-KEGG associations uniquely revealed by the addition of novoMP-DB, enabling the identification of 175 unique pathways for 48 species.

Our approach represents a novel orthogonal strategy for FDR control of novoMP-detected peptides, validating their confidence. The evaluation of CScore and Posterior Error Probability (PEP) showed that precursors from the two sources exhibited similar distributions (Figure 2C). Furthermore, novoMP-DB peptides demonstrated equal or slightly better quantitative precision compared to database-searched peptides across various sample loadings and gradient lengths (Supp. Figure 2B).

At the taxonomic level, novoMP-DB peptides contributed up to 92.9% of annotated taxa at different ranks (Figure 2D). Across the dataset, 825 species were annotated (using 3 species-specific peptides as cut-off), with novoMP-DB peptides enabling the detection of 631 (Figure 2E). This represents a 6 times increase in taxonomic coverage compared to our previous state-of-the-art DIA-PASEF^11^. Notably, 253 species would not have met the minimum cutoff of three species-specific peptides without the addition of novoMP-DB peptides, and 114 species were exclusively identified through unique novoMP-DB peptides (Figure 2E). From the 118,937 identified protein groups (PGs), 47,739 groups include proteins originating from novoMP-DB, among which 26,149 include proteins uniquely discovered by novoMP-DB (Supp. Figure 2C). These PGs spanned all 24 functional Clusters of Orthologous Genes (COG) categories, with minor differences in KEGG pathway counts (Supp. Figure 2D). Detailed functional analysis showed that proteins unique to novoMP-DB were largely represented in COG categories like RNA processing (COG-A), chromatin dynamics (COG-B), extracellular structures (COG-W), nuclear structure (COG-Y), and cytoskeleton (COG-Z) (Figure 2F). Overall, uMetaP enabled the study of 199 KEEG additional pathways compared to our previous work^11^. Examining species-to-function links, we demonstrated how de novoMP-DB proteins uniquely revealed 175 species-KEGG associations from 48 species (Figure 2G). Interestingly, uMetaP uncovered previously hidden functions by identifying 1,043 proteins (196 proteins originated from novoMP-DB) of unknown function^12^ (PUFs). Further, we identified 2,342 small proteins^13, 14^ (sProt; 321 proteins originated from novoMP-DB), and 581 proteins (86 proteins originated from novoMP-DB) with predicted antimicrobial peptide sequences^15^ (AMPs; Supp. Figure 2E). Altogether, uMetaP powered by DIA-PASEF considerably improves taxonomic and functional coverage, as well as quantification quality in complex metaproteomics samples, significantly benefiting the study of previously hidden host-microbial interactions.

### Redefining the detection limit of “dark metaproteome”

The human gut microbiome harbors an average of 200 bacterial species^19, 20^ comprising a core of abundant species present in most individuals^2, 3^ and a second pool formed by low-abundant species (more than 50% of the total), underlining the increasingly important inter-individual variability of microbiome profiles in health and disease. Current metaproteomic approaches do not achieve sufficient sensitivity to study these low-abundant species^4^. We hypothesized the benefits of uMetaP to significantly improve the study of this uncharacterized “dark” metaproteome.

We set to develop an approach to calculate the real lower limits of detection (LoD) and quantification (LoQ) by calculating the number of bacterial cells that can be accurately identified and quantified in a complex microbial sample. To minimize identification uncertainty, we used stable isotope labeling by amino acids in cell culture (SILAC) for *Ligilactobacillus murinus* (*L. murinus*), a bacterium native to the mouse gut microbiome. DDA-PASEF analysis of the SILAC culture confirmed an average incorporation efficiency of 97.42% (Supp. Figure 3A; Supp. Table 3). Additionally, we employed *Salinibacter ruber* (*S. ruber*) as an exogenous spiked bacteria (Figure 3A). We observed that the number of detected peptides and protein groups declined as we decreased the number of SILAC-labeled *L. murinus* and unlabeled *S. ruber* cells spiked into 10 mg of mouse feces (Figure 3A-B; Supp. Figure 3B). After applying strict filtering criteria for taxonomic identifications (including a non-spiked control; see Methods), we identified 6 and 20 peptides for *L. murinus* and *S. ruber*, respectively, when spiked 10,000 cells (Figure 3B; Supp. Table 4). Visual inspection of selected spectra confirmed these identifications (examples in Supp. Figure 3C-D). By extracting precursor ions and fragments from DIA-PASEF spectra, we determined a reliable LoQ of 1 million *L. murinus* cells and 5 million *S. ruber* cells (examples in Figure 3C-D). The differences in LoQ for each bacteria possibly reflect differences in bacterial size (Supp. Figure 3E) and protein content.

**Figure 3:**
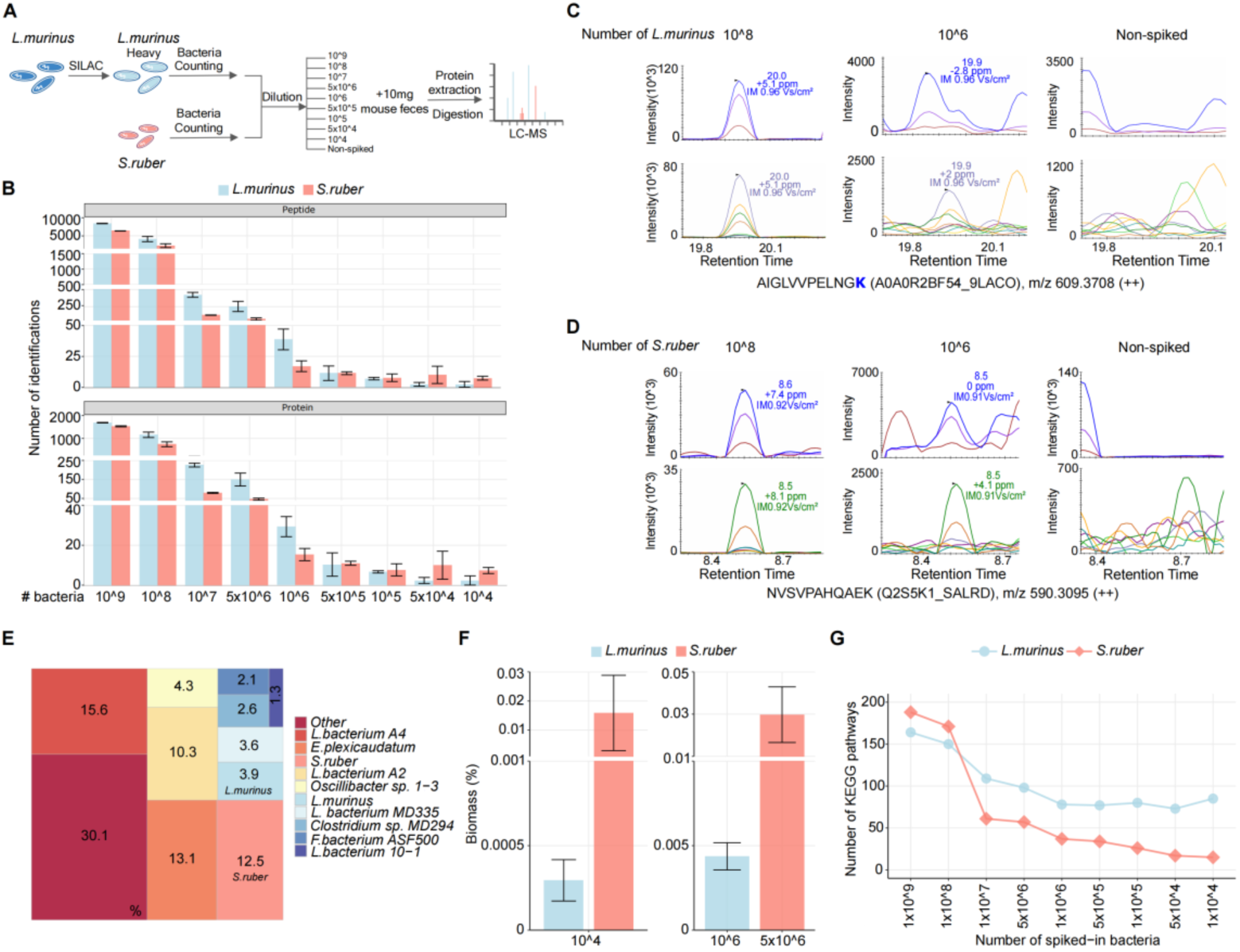
Redefining the detection limit of the “dark metaproteome” using uMetaP workflow. (A) Workflow combining SILAC labelling and spiking of *Ligilactobacillus murinus* (*L. murinu*s) and spiking of unlabeled exogenous *Salinibacter ruber* (*S. ruber*) into mouse feces. Each spike-in includes three replicates. (B) Number of peptides (top) and protein groups (bottom) identified for *L. murinus* and *S. ruber* across bacterial cell inputs. Error bars show the standard deviations across triplicates. (C-D) Representative extracted MS/MS spectra from complex DIA-PASEF data of peptides from *L. murinus* (C) and *S. ruber* (D) to illustrate reliable limit of quantification (LoQ), in comparison with the 10^^8^ spike-in and the non-spiked control. (E) Microbial community biomass composition (10^^8^ spike-in samples) showing that eight species (excluding spiked *L. murinus* and *S. ruber*) account for 53.5% of the total microbiota biomass. (F) Biomass percentages for spiked *L. murinus* and *S. ruber* cells at 10^4 (LoD), 10^6 (LoQ for *L. murinus*), and 5 x 10^6 (LoQ for *S. ruber*) input levels. (G) Number of KEGG pathways annotated from identified protein groups as bacterial inputs decreased.

Species abundance within a microbial community is an important parameter for microbiome studies. By summing the intensities of species-specific peptides, we showed that uMetaP abundance assessments are driven by a limited number of species, with just eight species (excluding spiked *L. murinus* and *S. ruber*) accounting for 53.5% of the microbiota biomass (Figure 3E; Supp. Table 5). Among 115,127 peptides identified in this dataset, 21,457 can be traced back to the novoMP-DB (Supp. Figure 3F), which contributed to the detection of species down to 0.006% relative abundance (data not shown). Peptide intensity analysis indicated that the 10,000 spiked *L. murinus* and *S. ruber* cells detected by uMetaP constituted 0.0003% and 0.0159% of the total biomass, respectively (Figure 3F; Supp. Table 5). Considering the spectral quality of precursor and fragment ions, we confidently quantified these spiked bacteria, representing 0.0044% for *L. murinus* and 0.0297% for *S. ruber* (Figure 3F). Based on genomic estimates (which assume 1 x 10^12^ bacterial cells per gram of mouse feces^21^), we achieved a LoD of 0.0001% (1 cell detected among 1 million) for both species, and an LoQ of 0.01% and 0.05% for *L. murinus* and *S. ruber*, respectively. These results significantly improved the previously reported limits in metaproteomics^4^ by up to 5,000-fold and are comparable to the deepest profiling efforts using full-length 16S rRNA^20^. Additionally, functional annotation of identified protein groups to KEGG pathways diminished below 100 million bacteria and plateaued at 1 million bacteria (Figure 3G). Remarkably, we annotated 85 and 18 functional pathways with as few as 10,000 *L. murinus* and *S. ruber* cells, encompassing a variety of metabolic and biosynthetic pathways.

Our data establish new detection and quantification limits in complex metaproteomic samples, enabling a more precise definition of individual functional microbiomes.

### Shedding light on microbial-metabolic circuits underlining tissue injury during intestinal inflammation *in vivo*

The mutual relationship between the microbiome and the host is essential for maintaining intestinal homeostasis, and its disruption plays a role in the onset and progression of diseases, including inflammatory bowel diseases (IBD)^22, 23^. It has been recently demonstrated how mitochondrial (MT) perturbation in intestinal epithelial cells (IECs) causes metabolic injury, a self-autonomous mechanism of tissue wounding associated with microbial dysbiosis^24^ and triggers the recurrence of chronic intestinal inflammation^25^.

Beyond taxonomic associations using shallow shotgun metagenomics, how metabolic changes in the intestinal epithelium select the growth of certain bacteria and how specific bacteria interfere with epithelial regeneration are unknown, which precludes understanding of host-microbiome interactions and defining potential therapeutical targets. We set uMetaP to the test by investigating the dynamics of microbial-host circuits in recurrent intestine inflammation *in vivo*.

To test the role of MT function in epithelial stem cell homeostasis, we took advantage of our published model of MT dysfunction in the intestinal epithelium, in which the MT chaperone heat shock protein 60 (Hsp60) is transiently deleted, specifically in mouse IECs (Figure 4A). This deletion triggered temporary mitochondrial dysfunction, leading to metabolic stress and transient structural changes in the colonic epithelium similar to the ones observed in patients suffering from intestinal inflammatory diseases^24^. We explored the microbiome shifts and the host functional changes during tissue injury by analyzing the colonic contents from control (Hsp60^fl/fl^) and metabolic injured (Hsp60^Δ/ΔIEC^) mice at two-time points after tamoxifen cessation: day 0 (D0) which corresponds to Hsp60 complete loss but there are no apparent histological aberrations, and D8 corresponding to the peak of metabolic injury^24^.

**Figure 4:**
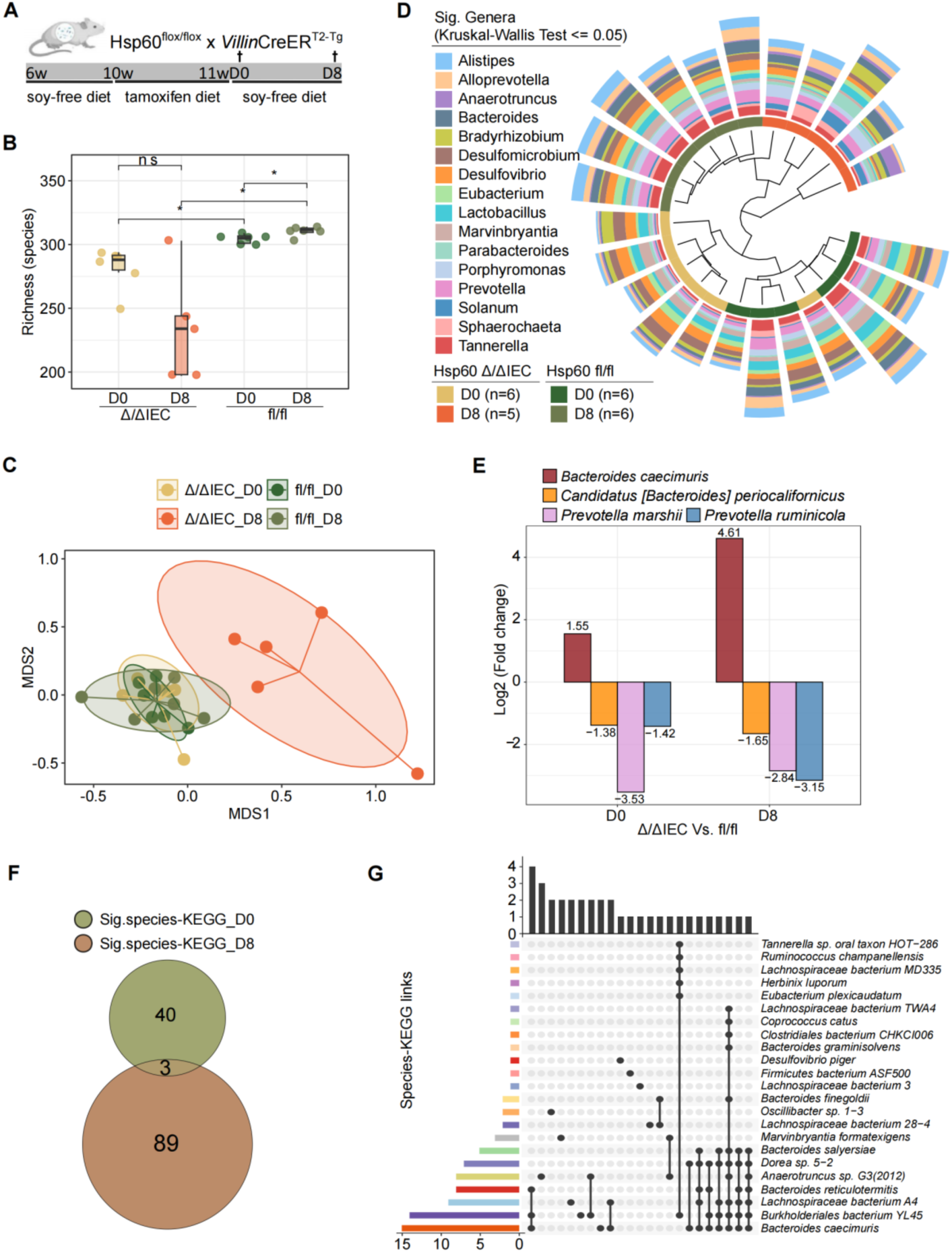
uMetaP sheds light on microbial-metabolic circuits underlining tissue injury during intestinal inflammation *in vivo*. (A) Schematic of the experimental design showing the strategy for the transient deletion of mitochondrial heat shock protein 60 (Hsp60) in mouse intestinal epithelial cells (IECs) using tamoxifen induction. Colonic contents from control (Hsp60^fl/fl^) and injured (Hsp60^Δ/ΔIEC^) mice were analyzed at day 0 (D0) and day 8 (D8) after the tamoxifen diet. (B) Microbial richness (alpha-diversity) in the 4 experimental groups. (C) β- diversity showing distinct microbial community structures between control and injured mice across D0 and D8. (D) Dendrogram shows the relative abundance of 16 genera significantly changed in response to metabolic injury discovered by uMetaP. The significance level was set with an adjusted p value ≤ 0.05 (Benjamini-Hochberg) after Kruskal-Wallis test. (E) Differential abundance of 4 commonly regulated species at D0 and D8. (F) Overlap of significantly regulated (T-test, adjusted p value ≤ 0.05) species-KEGG pathways at D0 and D8. (G) UpSet plot showing uniqueness and shareness of significantly regulated KEGG pathways among 23 species at D8.

Overall, peptide and protein identifications revealed distinct proteomic profiles between the two genotypes, with both host and microbiome IDs decreasing at D8 uniquely in the metabolic injured model (Supp. Figure 4A). uMetaP detected more than 300 species in all four experimental groups, significantly increasing the taxonomic coverage reached by 16S rRNA and shallow shotgun metagenomic sequencing on the same samples (Supp. Figure 4B). The metabolic injury phenotype was associated with significant changes in microbiota richness (Figure 4B) and community structure (β-diversity; Figure 4C). Differential abundance analysis identified 16 significantly altered genera across the four conditions, with 10 genera, including *Bacteroides*, *Anaerotruncus*, and *Parabacteroides*, showing increased abundance during injury, while *Lactobacillus*, for instance, decreased at D8 compared to the other groups (Figure 4D; Supp. Figure 4C), reflecting microbiome taxonomic adaptation to IEC dysfunction. At the species level, 25 and 18 differentially abundant species were observed at D0 and D8, respectively, in response to metabolic injury (fl/fl compared to Δ/ΔIEC; Supp. Figure 4D-4E). Four species were commonly regulated at both time points, with Bacteroides caecimuris uniquely showing increased abundance from D0 on already and more accentuated at D8 (Figure 4E), suggesting their potential role in modulating colon metabolic injury. Remarkably, the above-reported taxonomic changes discovered by uMetaP mirrored (e.g., β-diversity, an increase of *Bacteroides caecimuris*) and extended (previously unreported changes in several genera and species) the metagenomic findings on these same samples^24^.

Functional analysis indicated 43 and 92 significantly regulated species-KEGG pathways at D0 and D8, respectively, with only three pathways in common (Figure 4F and Supp. Table 6), underscoring unique functional shifts within the microbial community in response to injury. Notably, while *Bacteroides caecimuris* increased in abundance at both time points, it only showed significant KEGG pathway alterations at D8 (15 pathways; Figure 4G and Supp. Figure 4F). Detailed analysis revealed that *Bacteroides caecimuris* was the only species showing a significant alteration of two pathways at D8 - Carbon fixation by Calvin cycle and Biosynthesis of Ansamycins (Figure 4G and Supp. Table 6), offering mechanistic hints on how this bacterium might modulate colonic microenvironments to dominate over other taxa during metabolic injury^24^.

These data demonstrate the benefits of uMetaP, beyond genomic findings, to shed light on taxonomical and functional alterations underlying intestinal tissue injury *in vivo*.

### Defining the druggable gut metaproteome: Mining the host proteome offers potential therapeutic targets in human intestinal inflammatory diseases

Similar to the druggable genome in cancer research^26^, where genes are prioritized for therapeutic targeting, mining the gut metaproteome could allow researchers to identify proteins that influence host-microbiota interactions during disease states as prime candidates for therapeutic intervention, particularly in inflammatory bowel diseases. However, the concept of the druggable gut metaproteome remains unexplored. We set out a strategy to test the translational potential of uMetaP characterizing the proteome changes of the host during intestinal tissue injury and setting up an orthogonal inter-species validation strategy with Chron’s disease patient data.

Host analysis revealed a higher number of significantly regulated proteins in the metabolic injured model at D8 than at D0 (990 and 144, Figure 5A and Supp. Figure 5A, respectively). Consistent with the observed functional shifts in the microbial community (Figure 4F), there was minimal overlap in enriched host functions between D0 (Supp. Figure 5B) and D8 (Figure 5B, Supp. Table 7). In line with the tissue injury phenotype, at D8, we found an enrichment in functional pathways related to tissue homeostasis and epithelial development. Moreover, functions associated with cytokine regulation and the ERK1/2 cascade highlighted the pro-inflammatory environment occurring during tissue injury. We obtained orthogonal validation of the metaproteomic findings by comparison with transcriptome data obtained from mucosal biopsies of Crohn’s Disease patients^27^ (CD; 343 samples from 204 patients originating from inflamed ileum or M0I, and the non-inflamed ileal margin or M0M). We found 490 significantly regulated proteins at D8 presented in the list of regulated human transcripts (Supp. Table 8), where several proteins (e.g. RELA^28^, NOS2^29^, and ITGAM^30^) are reported to be linked to intestinal inflammatory diseases in humans.

**Figure 5:**
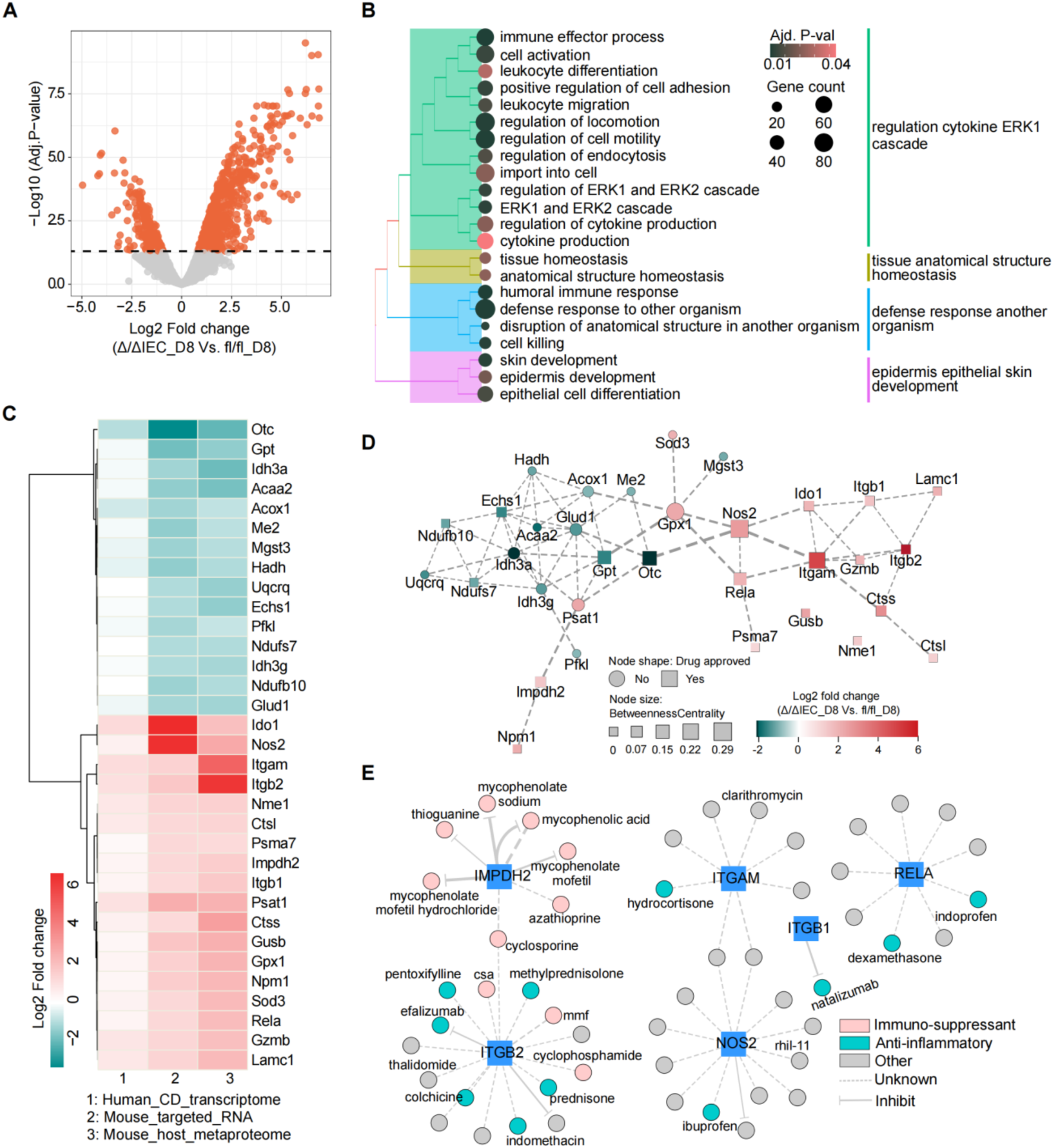
Mining host proteome changes during intestinal tissue injury defines the “druggable metaproteome” and reveals potential therapeutic targets for inflammatory diseases. (A) Volcano plot showing the log2 fold changes of significantly regulated proteins (Δ/ΔIEC_D8 vs. fl/fl_D8). (B) Functional enrichment analysis of significantly regulated proteins at D8 highlights pathways related to tissue homeostasis, epithelial development, cytokine regulation, and the pro-inflammatory ERK1/2 cascade. Gene counts and adjusted p-values are visualized by size and color, respectively. (C) Heatmap showing the directionality of log2 fold changes for 33 proteins consistently regulated in mouse metaproteomics (colonic contents), mouse targeted RNA analysis (colon tissue), and human transcriptomics datasets from Crohn’s patients (ileum biopsy). (D) Protein-protein interaction network analysis of 33 proteins. Node shapes indicate whether the target protein has approved drugs based on drug- gene interaction analysis. (E) Examples of drug-gene interaction network of ITGAM, ITGB2, RELA, IMPDH2, NOS2, and ITGB1. Node colors represent therapeutic categories (e.g., anti- inflammatory, immunosuppressant), and the edge types indicate the drug effects (inhibitory or unknown).

We set an integrative analysis workflow to prioritize interesting proteins, highlighting their biological importance, therapeutic potential, and available strategies for targeting intestinal inflammatory diseases. To further narrow the list of 490 regulated proteins, we detected 97 proteins represented among the list of transcripts expressed at D8 with tissue injury in another set of mouse colonic samples^24^ (Supp. Table 8). Finally, by selecting candidates with a similar directionality of regulation (up- or down-regulated) in all three datasets (mouse proteomics, human transcriptomics, and mouse transcriptomics), we limited the list to 33 proteins upon tissue injury (Figure 5C). Functional annotation analysis highlighted biological processes with roles in specific metabolic pathways, in molecular functions focused on oxidoreductase activity, and in cellular components related to the mitochondria (Supp. Figure 5C and Supp. Table 9). Protein interaction network analyses showed differential connectivity for down- and up-regulated proteins, with specific nodes playing key roles as "hubs" in the network controlling the flow of information during tissue injury, as shown by their higher betweenness centrality (Figure 5D). Notably, several proteins with the bigger centrality scores recapitulate known biology underlying inflammatory intestinal diseases in humans (e.g., Rela^28^, Nos2^29^, and Itgam^30^). We defined the druggable metaproteome by investigating drug-gene interactions, resulting in 204 interactions corresponding to 187 drugs (Supp. Table 10) for 20 out of the 33 genes/proteins. Interestingly, 77 of the drugs found are approved for several indications associated with human inflammatory diseases, including a current treatment for Crohn’s disease (e.g., Natalizumab^®^, targeting ITGB1; Figure 5E), an anti-inflammatory drug for IBD treatment (e.g., prednisone, targeting ITGB2; Figure 5E), and other target genes/proteins with high centrality in the interaction network (e.g., Hydrocortisone targeting ITGAM; Figure 5E). Moreover, we revealed another set of 57 drugs approved for other indications (Figure 5E; e.g., rhil-11 and clarithromycin), as well as 110 not approved compounds (Supp. Table 10).

Our data show the translational potential of uMetaP for hypothesis generation and prioritization of target proteins for review by clinical experts to aid in treatment decision- making.

## DISCUSSION

Metaproteomics holds significant potential to advance microbiome research. However, current methods struggle to achieve high sensitivity, deep protein profiling, and quantitative accuracy and precision. As a result, many medium- and low-abundance taxa identified by widely used genomic methods, along with their functional repertoires, are often not characterized. In uMetaP, we combined the latest LC-MS technology with a new de novo strategy to address these limitations. We demonstrated the benefits of uMetaP in meaningful biological scenarios using an *in vivo* mouse model of metabolic injury and by benchmarking our findings against human transcriptomic data from Crohn’s patients. The translational potential of our data was demonstrated through a detailed drug-gene analysis, enabling hypothesis-driven drug repurposing efforts.

In recent years, metaproteomics has advanced by integrating sophisticated mass spectrometry platforms^11, 31^, superior data acquisition methods (e.g., DIA-PASEF^11^), and machine-learning-based data analysis^11, 32^. Nonetheless, metaproteomic studies are still limited by database (DB) construction, with classic approaches relying on reference catalogues and database-search workflows, which capture only a small fraction of proteomic diversity within complex samples. Our data show that around 70% of the spectral information acquired by uMetaP was not utilized. Metaproteomics would greatly benefit from de novo sequencing solutions. However, the spectral complexity of these samples and the lack of methods for controlling de novo peptide confidence limit its application^16^.

We addressed this challenge by developing novoMP, a de novo strategy tailored for metaproteomics DB construction. Compared to previous studies^16^, novoMP is unique in three key aspects. First, it is the first de novo algorithm trained on the PASEF data structure, obtained from various timsTOF platforms, different species, and using different cleavage enzymes. Second, it implements a multi-layered quality control pipeline to select high-confidence de novo PSMs rigorously. Third, it offers a novel orthogonal FDR validation method using DIA-PASEF, demonstrating equivalent confidence in novoMP-DB peptides compared to peptides obtained through classical database search workflows.

We significantly expanded the taxonomic and functional representation in metaproteomic databases by combining the depth, sensitivity, and spectral quality of uMetaP with novoMP. Interestingly, the DIA-PASEF analysis with solely novoMP-DB covers more than 99% of the COG and KKEG pathways (Supp. Figure 2D), strongly suggesting the possibility of reaching maximum functional coverage without the need for metaproteomic DB construction in future studies. Moreover, the benefits of novoMP in our previously published DDA-PASEF data could be extended to PASEF datasets acquired in previous metaproteomic studies^32, 33, 34^. We provide the metaproteomic community with a roadmap for increasing confidence in de novo solutions and offer the most extensive mouse metaproteomic DB composed of 208,254 proteins, representing 774 microbial species and 447 KEGG pathways.

Combined with DIA-PASEF, uMetaP surpasses current proteotyping standards^4^, the most optimistic performance forecast in the field^35^, and a preliminary evaluation of the Orbitrap Astral mass spectrometer^31^. The fast analysis times and the low variability reached make uMetaP a promising tool for large-scale metaproteomic studies. Integrating novoMP into DB construction enabled the identification of an additional 28% of proteins, and 80% of taxa (Supp. Figure 2C; Figure 2E). Beyond identification, uMetaP demonstrates exceptional precision, reliability in quantification, and ultra-high sensitivity. These benefits were demonstrated by establishing the first reliable LLoD and LLoQ in a complex metaproteome. Unlike previous approaches^4^, we accounted for sample preparation losses by using a SILAC-labeled bacterium (*L. murinus*) and an exogenous bacterium (*S. ruber*). As a result, uMetaP can detect a single bacterium in a theoretical background of 1 million, representing a 5,000- fold improvement^4^. Importantly, MS2 spectra showed a shift in the reliable quantification limit for *L. murinus* and *S. ruber*, which is likely applicable across metaproteomes and highlights the importance of rigorous spectral quality control for accurate peptide quantification. Establishing LLoD and LLoQ for the gut "dark metaproteome" has important implications. By lowering the thresholds for reliably identifying and quantifying bacterial species and their protein products, researchers can better capture the functional contributions of often-overlooked low-abundance species. This is critical for fields that require ultra-sensitivity - from marine metaproteomics^36^ to clinical metaproteomics, where subtle but clinically important changes in pathogenic microorganisms demand early detection. Moreover, reliable quantification of medium- and low-abundance species will help answer key questions about individualized and healthy microbiome profiles^2, 3^.

Our results on a transgenic mouse model of colonic tissue injury demonstrated the potential of uMetaP for discovering novel host-microbiome interactions in a relevant *in vivo* context. In addition to mirroring the taxonomic findings reported using genomic methods^24^, uMetaP offers greater sensitivity, allowing earlier detection of taxonomic and functional alterations underlying causal disease mechanisms. Urbauer et al demonstrated that *Bacteroides caecimuris* increases in abundance during metabolic injury at day 8 after the start of tissue injury and that mono-colonization of germ-free Hsp60 knock-out mice with *B. caecimuris* is sufficient to recapitulate the disease phenotype. Similarly, our data showed an increase in *Bacteroides caecimuris* at day 8. Notably, uMetaP also detected this increase during the first 24 hours after tamoxifen cessation. This is a significant improvement in the temporal sensitivity for detecting taxonomic changes compared to genomic and immunohistochemistry methods. Moreover, we extended the previously known dysbiosis signature by detecting significant abundance changes in additional bacterial species.

Bacteroides species are known to adapt well to inflammatory and stressed conditions^37^, potentially explaining the observed selective expansion in response to colonic injury. However, the mechanisms leading to this selective advantage remain unknown. Our functional data provide plausible mechanisms by detailing metabolic reprogramming during disease progression *in vivo*. We identified *B. caecimuris* as the bacterial species with the most KEGG pathways altered at day 8. Notably, two altered KEGG pathways were unique to this species: carbon fixation by the Calvin cycle and biosynthesis of ansamycins. The simultaneous upregulation of these pathways may provide a competitive advantage in the gut microbiome for *B. caecimuris*, especially in the context of mitochondrial dysfunction in the intestinal epithelium. On the one hand, the impaired mitochondrial function caused by the Hsp60 mutation^38^ could lead to reduced CO₂ production due to decreased TCA cycle activity. Upregulating carbon fixation via the Calvin cycle equips the bacteria with greater metabolic flexibility, allowing them to utilize even small amounts of CO₂, which could provide an advantage over competitors. On the other hand, ansamycins, such as rifamycins, are antibiotics produced by certain bacteria^39^. In the disrupted gut environment caused by Hsp60 deletion, the selective elimination of sensitive competitors could allow the ansamycin- producing bacteria to dominate. Interestingly, Bacteroides species are not primary producers of ansamycins^40^. Our data discovered specific metabolic adaptation by *Bacteroides caecimuris*, potentially contributing to its expansion during tissue injury as detected by metagenomics^24^ and uMetaP.

Beyond classical functional analysis of host proteins, we explored the translational potential of our findings. We introduced and explored the concept of a “druggable metaproteome”: The collection of host and microbiota proteins within a given environment that possess the structural and functional properties necessary to be targeted by pharmaceutical agents. This concept supports drug discovery and repurposing efforts. The orthogonal inter-species validation with transcriptomic data from Crohn’s patient biopsies^27^ validated changes in over 400 mouse proteins. This underscores the unique strength of metaproteomics as an -omic technique for detecting functional changes in host physiology. To prioritize proteins for future studies and identify potential therapeutic strategies for intestinal inflammatory diseases, we combined functional, molecular network, and drug-gene interaction analyses. We identified more than 200 potential drug-protein interactions, including immune-suppressants used in Crohn’s disease (e.g., natalizumab), anti-inflammatory drugs for IBD treatment (e.g., prednisone), and approved drugs for other uses. Follow-up studies using pre-clinical mouse models or human volunteers are needed to test data-driven hypotheses suggesting specific drug repurposing or combinatorial treatments.

By integrating cutting-edge LC-MS technology, developing a novel de novo strategy, testing these advancements on an *in vivo* disease model, and introducing the concept of the “druggable metaproteome,” our study advances metaproteomics, highlighting its potential in microbiome research, particularly in unraveling host-microbiome interactions and their crucial roles in health and disease.

## MATERIAL AND METHODS

### Reagents

All reagents were purchased from Sigma-Aldrich (St. Louis, Missouri) if not mentioned otherwise. Acetonitrile (ACN) and formic acid (FA) were purchased from Fisher Scientific (Hampton, New Hampshire; both FA and ACN were liquid chromatography-mass spectrometry (LC-MS) grade). LC-MS grade water from Sigma-Aldrich was used for all solutions. Protease inhibitor (Complete Ultra Tablets Mini) was purchased from Roche, Basel, Switzerland.

### Animals and housing conditions

In-house bred C57BL/6J mice were used for data presented in Figures 1-3. Housing and operation of mice were carried out with the approval of the University of Vienna animal care and use committee (license number 2021-0.138.925). All mice used in this study were group-housed in individually-ventilated cages in a 12-hour light/dark cycle in the animal facility with water and food *ad libitum*.

Mice used in the *in vivo* experiments (Figure 4 and Figure 5) were only male animals. Details of the animal models can be found in our previous study^24^. Briefly, Hsp60^flox/flox^ mice and Hsp60^flox/flox^ x VillinCreER^T2-Tg^ mice were generated as described previously^41^ to create IEC-specific Hsp60 knockout mice via tamoxifen induction (Hsp60^Δ/ΔIEC^). For conditional Hsp60 deletion, Hsp60flox/flox x VillinCreERT2-Tg mice and appropriate control mice were kept on phytoestrogen-reduced diet 1005 (V1154-300, Ssniff) for four weeks under SPF conditions. Afterwards, mice received 400mg tamoxifen citrate per kg chow feed (CreActive T400 (10mm, Rad), Genobios) ad libitum for 7 days. After the induction phase, tamoxifen diet was replaced with the phytoestrogen-reduced diet. During and after the induction phase, mice were monitored daily and aborted when a combined score considering weight loss, changes in stool consistency, general behavior, and general state of health was reached. Animals were sacrificed at the indicated time points. All mice and their respective genotypes were generated and maintained on an in-house crossing of C57Bl/6N and C57Bl/6J background. All mice were housed under specific pathogen-free (SPF) conditions according to the criteria of the Federation for Laboratory Animal Science Associations (FELASA) (12-hour light/dark cycles at 24–26°C) in the mouse facility at the Technical University of Munich (School of Life Sciences Weihenstephan). All mice received a standard diet (autoclaved V1124-300, Ssniff) *ad libitum*, autoclaved water and were sacrificed by CO_2_ or isoflurane.

### Protein extraction and SP3-assisted protein digestion for metaproteomics analysis

The procedures from fecal collection to final peptide preparation were performed as previously described^11^.

### High pH reversed-phase fractionation of pooled peptides

The peptide fractionation kit was purchased from Fisher Scientific (Cat. 84868). A total of 40 µg pooled fecal peptides were processed according manufactures instruction. Eight peptide factions were dried using vacuum centrifugation and then re-suspended in 30 µL of MS-grade water. The peptide concentration was measured in duplicate using NanoPhotometer N60 (Implen, Munich, Germany) at 205 nm. Peptide samples were acidified with formic acid to a final concentration of 0.1% and were stored at -20°C until LC-MS/MS analysis.

### Liquid chromatography-mass spectrometry configurations

Nanoflow reversed-phase liquid chromatography (Nano-RPLC) was performed on NanoElute1 and NanoElute2 systems (Bruker Daltonik, Bremen, Germany) coupled with timsTOF Pro and timsTOF Ultra (Bruker Daltonik, Bremen, Germany) via CaptiveSpray ion source, respectively. Mobile solvent A consisted of 100% water containing 0.1% FA and mobile phase B of 100% acetonitrile containing 0.1% FA.

### Data dependent acquisition (DDA-PASEF) of fractionated peptides on timsTOF Ultra and timsTOF Pro

Twenty-nanograms of each peptide fraction were loaded on an Aurora^TM^ ULTIMATE column (25 cm x 75 µm) packed with 1.6 µm C18 particles (IonOpticks, Fitzroy, Australia) with a total gradient time of 66 minutes. The mobile phase B was linearly increased from 5 to 23% in 56 minutes with a flowrate of 0.25 µL/min, followed by another linear increase to 35% within 4 minutes and a steep increase to 90% in 1 minute. The mobile phase B was maintained at 90% for the last 5 minutes with a flowrate increase from 0.25 µL/min to 0.35 µL/min. On both timsTOF Ultra and timsTOF Pro, the TIMS analyzer was operated in a 100% duty cycle with equal accumulation and ramp times of 166 ms each. Specifically, 5 PASEF scans were set per acquisition cycle (cycle time of 1.03 s) with ion mobility range from 0.7 to 1.3 (1/k0). The target intensity and intensity threshold were set to 14000 and 500 respectively. Dynamic exclusion was applied for 0.4 minutes. Ions with m/z between 100 and 1700 were recorded in the mass spectrum. Collision energies were dependent on ion mobility values with a linear increase in collision energy from 1/K0 = 0.6 Vs/cm² at 20 eV to 1/K0 = 1.6 Vs/cm² at 59 eV. The TIMS analyzer was operated in 100% duty cycle with 100 ms defined for accumulation and ramp time. Ten PASEF scans were set per acquisition cycle (cycle time of 1.17 s) with ion mobility range from 0.65 to 1.45 (1/k0). The target intensity and intensity threshold were set to 10000 and 1750 respectively.

### Data independent acquisition (DIA-PASEF) on timsTOF Ultra

Peptides were loaded onto an Aurora^TM^ ULTIMATE column (25 cm x 75 µm) packed with 1.6 µm C18 particles (IonOpticks, Fitzroy, Australia) with a total gradient time of either 30 minutes or 66 minutes on a NanoElute2 system in triplicates. In the 30-min separation, the mobile phase B was linearly increased from 5 to 23% in 18 minutes with a flowrate of 0.25 µL/min, followed by another linear increase to 35% within 4 minutes and a steep increase to 90% in 2 minutes. The mobile phase B was maintained at 90% for the last 4 minutes with a flowrate increase from 0.25 µL/min to 0.35 µL/min. The composition of mobile phase B over the 66- min separation was the same as described above for the fractionated peptide samples. For the results presented in Figure 2, precursors with m/z between 400 and 1000 were defined in 8 scans (3 quadrupole switches per scan) containing 24 ion mobility steps in an ion mobility range of 0.64 – 1.45 (1/k0) with fixed isolation window of 25 Th in each step. The acquisition time of each DIA-PASEF scan was set to 100 ms, which led to a total cycle time of around 0.95 sec. For results presented in Figure 3, 25ng peptides were separated on the NanoElute 2 system with a 30-min gradient. Precursors with m/z between 350 and 800 were defined in 6 scans (3 quadrupole switches per scan) containing 18 ion mobility steps in an ion mobility range of 0.64 – 1.2 (1/k0) with fixed isolation window of 25 Th in each step. The acquisition time of each DIA-PASEF scan was set to 100 ms, which led to a total cycle time of around 0.74 sec. For data presented in Figure4-5, 50 ng peptides were separated on the NanoElute 2 system with a 66-min gradient. Precursors with m/z between 350 and 1150 were defined in 13 scans containing 32 ion mobility steps in an ion mobility range of 0.65 – 1.35 (1/k0) with fixed isolation window of 25 Th in each step. The acquisition time of each DIA-PASEF scan was set to 100 ms, which led to a total cycle time of around 1.48 sec.

### DDA-PASEF data processing

Fractionated data generated using timsTOF Ultra and timsTOF Pro were separately submitted to MSfragger^42^ (Version 4.0) integrated in FragPipe computational platform (Version 21.1), searching against the MGnify mouse gut protein catalogue v1.0 (https://www.ebi.ac.uk/metagenomics/genome-catalogues/mouse-gut-v1-0, referred as PD1). The decoy database was generated with reversed sequences. Trypsin was specified with a maximum of two missed cleavages allowed. The search included variable modifications of methionine oxidation and N-terminal acetylation and a fixed modification of carbamidomethyl on cysteine. The mass tolerances of 10ppm and 20 ppm were set for precursor and fragment, respectively. Peptide length was set to 7 to 50 amino acids with a mass range from 500 to 5000 Da. The remaining parameters were kept as default settings. During the validation, MSBooster (Version 1.1.28) was used for rescoring and Percolator^43^ (version 3.6.4, default parameters) was used for PSM validation. FDR level was set to 1% for PSM, peptide and protein. The identified proteins from the search formed a sample-specific protein database (PD2) containing 53,502 protein sequences. For assessing the labelling efficiency of *L. murinus*, the data was searched against the standard proteome of *L. murinus* downloaded from Uniprot (PD3, https://www.uniprot.org/proteomes/UP000051612, accessed on 2023-07-19) containing 1,971 protein sequences. The rest parameters were kept the same in MSfragger as aforementioned.

### Bacterial culture of *L. murinus* and *S. ruber*

*Ligillacotobacillus murinus* (DSM 20452, *L. murinus*) and *Salinibacter ruber* (DSM 13855, *S. ruber*) were purchased from DSMZ (Braunschweig, Germany). All culture media were autoclaved right after the preparation. *L. murinus* was activated in 5 mL MRS medium (CARLROTH, Karlsruhe, Germany; prepared according to the manufacturer’s instructions) and incubated for 24 hours at 37 °C with 220 rpm agitation. At the end of this incubation period, 1 mL of the *L. murinus* culture was taken and centrifuged at 3200 g for 5 minutes at 4 °C. The supernatant was carefully removed, and the bacterial pellet was gently resuspended in 5 mL SILAC-heavy medium (Glucose 10 g/L, KH_2_PO_4_ 3 g/L, K_2_HPO_4_ 3 g/L, sodium acetate 5 g/L, ammonium citrate dibasic 1 g/L, MgSO_4_·7H_2_O 0.2 g/L, MnSO_4_·4H_2_O 0.05 g/L, Tween-80 1 g/L, L-alanine 0.05 g/L, L-arginine-HCl (13C_6_, 15N_4_; Fischer Scientific) 0.05 g/L, L-asparagine 0.1 g/L, L-aspartic acid 0.1 g/L, L-cysteine 0.2 g/L, L-glutamine 0.1 g/L, L-glutamic acid 0.1 g/L, glycine 0.05 g/L, L-histidine 0.05 g/L, L-isoleucine 0.05 g/L, L-leucine 0.05 g/L, L-lysine-2HCl (13C_6_, 15N_2_; Fischer Scientific) 0.05 g/L, L-methionine 0.05 g/L, L-phenylalanine 0.05 g/L, L-proline 0.05 g/L, L-serine 0.05 g/L, L-threonine 0.05 g/L, L-tryptophan 0.05 g/L, L-tyrosine 0.05 g/L, L-valine 0.05 g/L, uracil 0.01 g/L, guanine 0.01 g/L, adenine 0.01 g/L, xanthine 0.01 g/L, biotin 0.01 g/, Vitamin Solution 2% (v/v)). The heavy-medium culture was incubated at 37 °C with 220 rpm agitation for 24 hours. Bacterial growth was monitored with spectrophotometric measurements (Eppendorf, Hamburg, Germany) at an optical density of 600 nm (OD600). An OD600 above 0.8 was aimed to ensure suitable growth conditions. For daily passage, 500 microliters of *L. murinus* culture were taken and transferred to another 5 mL SILAC-heavy medium. The labelling efficiency was evaluated on timsTOF Pro after 10 passages in heavy-medium culture. *S. ruber* was cultured in 5 mL DMSZ-936 medium according to the recommendation (https://mediadive.dsmz.de/medium/936) at 37 °C with 220 rpm agitation. The duration between passages for *S. ruber* was around 7 days due to its slow growth. For enlarged culture, 1 mL/each of *L. murinus* and *S. ruber* cultures were transferred to 30 mL mediums, respectively. At the end of cultivation, 2 mL bacteria aliquots were made and pelleted at 3200 g for 5 minutes at 4 °C, and one of the aliquots was resuspended in 2 mL of either pre-chilled PBS (*L. murinus*) or DSMZ-936 medium (*S. ruber*). The resuspended bacteria were further serial diluted (2-50 times dilution) with either PBS (*L. murinus*) or DSMZ-936 medium (*S. ruber*) for bacteria counting using QUANTOM Tx Microbial Cell Counter (BioCat, Heidelberg, Germany) according to the procedures supplied with the device. The rest of the aliquots were snap-frozen in liquid nitrogen and stored at -80°C until further use.

### *L. murinus* and *S. ruber* Spike-in experiment

Counted *L. murinus* and *S. ruber* stocks were resuspended and diluted in pre-chilled PBS to reach various numbers (ranging from 1 x 10^4^ to 1 x 10^9^) in triplicates. The same number of *L. murinus* and *S. ruber* were mixed with 10 mg of mouse feces and subjected to protein extraction together (as previously described^11^). To ensure a consistent spike-in background, the fecal sample used here was collected and pooled from the same mouse in two consecutive days at the same hour. The resulting peptide samples were analyzed on the timsTOF Ultra in a 30-min gradient as described above with 25 ng of peptide per sample. The workflow is illustrated in Figure 3A.

### Labelling efficiency check for *L. murinus*

The labeling efficiency was checked by analyzing the heavy-labeled culture of *L. murinus* in DDA-PASEF mode, and the data were searched against its reference proteome (PD3, https://www.uniprot.org/proteomes/UP000051612, accessed on 2023-07-19) in Fragpipe with arginine (+10) and lysine (+8) as additional variable modifications. As a result, a total of 60,485 PSMs were identified (1% FDR), corresponding to 12,852 unique stripped peptide sequences were identified. In cases where multiple PSMs were assigned to the same peptide, only the most intense PSMs of one peptide was kept for both labeled and non-labeled forms if the latter was co-identified. If the peptide was identified only in either the heavy- or light-labeled form, the missing intensities were assigned a value of 1 to apply the following formula (doi:/10.1016/j.jprot.2018.12.025) for each peptide to calculate the labelling efficiency: Peptide labeling efficiency = (Intensity_Heavy / (Intensity_Heavy + Intensity_Light)) x 100. The average of calculated efficiency (97.42%) for all peptides was presented in the study.

### Training of Novor algorithm with PASEF datasets and performance evaluation

In order to obtain a robust tool for de novo sequencing using 4-dimention PASEF data, a custom version of Novor^18^ (BPS-Novor) was generated by training Novor’s decision tree-based scoring functions on over 1, 750,000 PSMs acquired in PASEF mode from a variety of timsTOF instruments. This training dataset included experiments with fixed collision energy measurements of deeply fractionated (a total of 60 high-pH offline fractions) peptide samples digested with GluC, Pepsin, Elastase, Chymotrypsin, and Trypsin. The ground truth data was taken from ProLuCID-GPU^44^ database search results filtered with 1% FDR with DTASelect^45^ at PSM level.

To evaluate the performance of the newly trained BPS-Novor, a publicly available mixed species (*H. sapiens, Yeast, E.coli*) dataset^46^ (ProteomeXchange ID: PXD014777) excluded in the training phase was used to determine the accuracy of the model. In addition, the performance of BPS-Novor was validated against K562 cell lysates digested with non-tryptic enzymes, specifically Elastase, Pepsin, GluC, and Chymotrypsin to ensure accuracy with mimicked non proteotypic peptides. These samples were analyzed on a 35 minute gradient using an EASY-nLC (Thermo Fisher) and a timsTOF Pro instrument. The precision and recall values were calculated as previously described.

### de novo sequencing of DDA-PASEF data

Fractionated data were submitted to BPS-Novor intergrated in ProteoScape (Bruker Daltonik, Bremen, Germany) for de novo sequencing. The mass tolerances for precursors and fragments were set to 20 ppm and 0.02 Da, respectively. Tryptic peptides with a maximum of two missed cleavages were allowed. Carbamidomethyl was set as a fixed modification on cysteine, and methionine oxidation and N-terminal acetylation were set as variable modifications. A maximum of two variable modifications per peptide was allowed. In addition, only the top candidate sequence per spectrum was exported in the output.

### Multi-tier filtering of de novo sequenced PSMs

The de novo sequencing outputs were imported into R and subjected to the following six filters sequentially. 1) De novo score: The first filter was based on the Novor de novo sequencing software, applying a score threshold of 65. 2) Charge state: We excluded PSMs with a charge state of 1 due to their less reliable fragmentation patterns. 3) Peptide length: we removed peptides shorter than seven amino acids to reduce the risk of ambiguous matches. 4) Mass error: We evaluated the mass error of sequenced precursors and retained only 95% of the sequenced PSMs that fell within the upper and lower cut-offs calculated using qnorm function in R based on the mass error distribution. 5) Retention time shift: Retention time predictions were performed using DeepLC^47^ (v2.2.27). We retained 95% of the remaining PSMs, which showed a strong correlation between observed and predicted retention times, based on the upper and lower cutoffs calculated using the qnorm function in R. 6) Collisional cross-section (CCS) shift: CCS predictions were performed using IM2Deep^48^ (v0.1.7). We retained 95% of the remaining PSMs that showed a strong correlation between measured and predicted CCS values, using cutoffs calculated as described above.

### Blast homology search of de novo sequenced peptides for the construction of microbial protein database

Unique peptides remaining after multi-tier filtering were subjected to a BLAST+ homology search^49^ to retrieve potential protein sequences for microbial protein database construction. The blastp function embedded in Diamond^50^ (v2.1.9; command line) was used to search against the non-redundant protein sequence database “nr.gz” (ftp://ftp.ncbi.nlm.nih.gov/blast, updated 2024-02-27). The search of de novo sequenced peptides in ultra sensitive mode was restricted to the following taxa due to the nature of our samples: bacteria (taxaID: 2), fungi (taxaID: 4751), archaea (taxaID: 2157), and viruses (taxaID: 10239). All BLAST searches used the PAM30 scoring matrix. The top 5 protein assignments per query sequence were listed in the output file (output format: 6). In addition, another search with same parameters but different output format (output format: 102) was performed to generate taxonomic classifications of sequenced peptides based on the lowest common ancestor (LCA) algorithm. To select the only one protein assignment per query sequence among the top 5 candidates, we used LCA-guided procedure. Specifically, if the taxonomic annotation of one protein candidate matches exactly the taxonomy assignment in the LCA output, then this candidate is kept. In the case that the taxonomic annotation of the protein candidates do not match exactly to the LCA output but belong to taxon rank in the LCA output, these candidates were kept. Finally, if the above two steps did not generate one protein per query sequence, the blast parameters (Bitscore, pident and e-value) will be applied to keep the most confident candidates. To further increase the quality of the blast search result, we applied a minimum of 80% cut-off for sequence identity, then further retrieved the protein sequences from NCBI using the protein sequence IDs in the blast output to form a microbial database based on novoMP (novoMP-DB; PD4). As a comparison, peptides identified using the aforementioned MSFragger search were subjected to the same blast homology search referred as DB-search (PD5) in the manuscript.

### DIA-PASEF data processing

DIA-NN^51^ (version 1.9) was used to process DIA-PASEF data in library-free mode to generate the predicted spectrum library. A deep learning-based method was used to predict theoretical peptide spectra along with their retention time and ion mobility. Trypsin/P was used for in silico digestion with an allowance of a maximum of 2 missed cleavages. Variable modifications on peptides were set to N-term methionine excision, methionine oxidation and N-terminal acetylation, while carbamidomethylation on cysteine was a fixed modification. The maximum number of variable modifications on a peptide was set to 2. Peptide length for the search ranged from 7 to 30 amino acids. The m/z ranges were specified accordingly depending on the experiment which aligned with the DIA-PASEF acquisition method, and fragment ions were set to a range from 100 to 1700. Mass accuracy for both MS1 and MS2 was set to automatic determination. Protein inference was set to “Protein names (from FASTA)” and the option of “Heuristic protein inference” was unchecked. Match-between-run (MBR) was enabled for cross-run analysis. RT-dependent cross-run normalization and QuantUMS^52^ (high precision) options were selected for quantification.

Generally, all searches in DIA-NN included a *Mus musculus* reference proteome (https://www.uniprot.org/proteomes/UP000000589, accessed on 2023.04.07) together with different microbial databases. Specifically, results presented in Figure2 and Figure4 eres searched against PD2, PD4 and PD5 (de-duplicated). Data shown in Figure3 was searched against PD2, PD4, PD5, as well as the standard proteome of *L. murinus* (PD3) and *S. ruber* (PD6; https://www.uniprot.org/proteomes/UP000008674, accessed on 2023-07-19). In addition to the searching parameters mentioned above, heavy isotopic labelling of arginine (+ 10.0082699 Da) and lysine (+8.014199 Da) were set as variable modifications.

The DIA-NN search outputs were further processed with the R package, DIA-NN (https://github.com/vdemichev/diann-rpackage), to calculate the MaxLFQ^53^ quantitative intensities for all identified peptides and protein groups with q-value ≤ 0.01 as criteria at precursor and protein group levels.

### DIA-PASEF spectrum visualization

Skyline^54^ (version 23.1.0.380) was used to visualize the spectra of peptides identified by DIA-NN. Briefly, the spectral library generated by DIA-NN after database searching was imported into Skyline to construct a library containing precursor information for the detected peptides. Precursors listed in the library and their associated fragment ions were then extracted from the raw DIA-PASEF data. During extraction, mass accuracy was set to 10 ppm for both precursors and fragments. To minimize false matches, only scans within 5 minutes of the retention times listed in the library were extracted.

### Taxonomic and functional annotation and quantification

iMetaLab^55^ (Version 2.3.0) was used for taxonomic annotation. Peptide sequences and their corresponding intensity data were imported into iMetaLab, and the built-in taxonomy database was used for mapping, with blanks ignored below the rank of Superkingdom and a minimum unique peptide count of 3 required. For the quantification of specific taxonomic ranks^11^, the annotation output was processed in R to extract peptides commonly detected across samples for taxonomic rank of interest (e.g genus, species). The intensity of each taxon was calculated by summing up the intensities of common peptides in each sample. The resulting summed intensities were log2-tranformed for statistical analysis.

The microbial protein databases used in this manuscript were annotated using EggNOG- mapper^56^ (http://eggnog-mapper.embl.de/) with default settings to retrieve potential functions and pathways.

### Taxon-specific functions analysis

Meta4P^57^ was used to analyze taxon-specific functions. The peptide quantification data from DIA-NN, taxonomic annotation output from iMetaLab, and functional annotation files from EggNOG-mapper were used as inputs for Meta4P. Quantification of taxon-specific functions was performed by summing the peptide intensities associated with specific functions. The resulting summed intensities were log2-transformed for statistical analysis.

### Identification filters of *L. murinus* and *S. ruber* for spike-in experiment

Peptide and protein identifications generated from DIA-NN search of the spike-in experiment were further filtered to ensure species-specific identifications: 1) Only heavy-labeled peptides were considered for *L. murinus* to exclude the interference from endogenous species. Heavy-labeled peptides assigned to *S. ruber* were removed as they represent false-matches. 2) Co-assigned peptides and protein groups shared between *L. murinus* and *S. ruber* were excluded. 3) Peptides assigned to *L. murinus* or *S. ruber* that were also identified in any of the non-spike controls (three replicates) were removed.

### Functional enrichment analysis of differentially expressed host proteins

Quantified host proteins were statistically compared in R using the ProTIGY package (https://github.com/broadinstitute/protigy) with a two-sample moderated t-test. Functional enrichment of differentially expressed host proteins was performed using the clusterProfiler^58^ R package, with all identified proteins in the study as background genes for enrichment analysis against the Gene-Ontology Biological Process database. The Benjamini-Hochberg method was used to adjust p values, with an adjusted p-value cutoff of 0.05 used to identify significantly enriched pathways.

Protein-protein interaction networks were analyzed using STRING within Cytoscape (v 3.10.2) under default parameters. Drug-gene interactions were retrieved using DGIdb^59^ (v 5.0.7) with default settings.

### Statistical analysis

The Kruskal-Wallis test was performed in R to identify significant differences in genera among conditions. Differentially expressed species and taxon-specific functions were analyzed using the limma package in R for the respective comparisons. The Benjamini-Hochberg method was applied for multiple comparisons in all statistical analyses.

## AUTHOR CONTRIBUTIONS

Conceptualization: D.G.V.; Experimental design: D.G.V., F.X., D.A., E.U., and D.H.; Biochemistry and mass spectrometry: F.X. and C.K.; Sample collection and preparation: F.X., M.B., R.K., E.U., and D.A.; Data analysis: F.X., D.G.V., J.K., T.S., Q.L., A.B., and B.M.; Writing: D.G.V., F.X., D.A. All authors edited and approved the final manuscript; study supervision: D.G.V.; project administration: M.S. and D.G.V.; Funding acquisition: M.S. and D.G.V.

## NOTES

M.S. received research awards and travel support from the German Pain Society (DGSS) both of which were sponsored by Astellas Pharma GmbH (Germany). MS received research awards from the Austrian Pain Society. MS received a one-time consulting honorarium from Grunenthal GmbH (Germany). None of these sources influenced the content of this study, and MS declares no conflict of interest. D.G.V. and M.S. have an ongoing scientific collaboration with Bruker (Center of Excellence for Metaproteomics University of Vienna - Bruker), however, this collaboration did not influence the content of the manuscript. D.G.V, F.X., M.B, R.K., A.B., D.A., D.H., and M.S. declare that they have no conflicts of interest. C.K., J.K., and T.S. are employees of Bruker Daltonics GmbH & Co. Q.L. and B.M. are employees of Rapid Novor.

## ETHICS

All animal experiments carried out at the University of Vienna were in strict accordance with institutional IACUC guidelines, international ARRIVE guidelines, and the principles of the 3Rs of animal research. All animal experiments carried out at the Technical University of Munich, as well as maintenance and breeding of mouse lines, were approved by the Committee on Animal Health Care and Use of the state of Upper Bavaria (Regierung von Oberbayern; AZ ROB-55.2-2532.Vet_02-14-217, AZ ROB-55.2-2532.Vet_02-20-58, AZ ROB-55.2-2532.Vet_02-18-37) and performed in strict compliance with the EEC recommendations for the care and use of laboratory animals (European Communities Council Directive of November 24, 1986 (86/609/EEC)).

## DATA AVAILABILITY

The mass spectrometry proteomics data have been deposited to the ProteomeXchange Consortium via the PRIDE partner repository with the dataset identifier PXD051792.

## MATERIALS AND CORRESPONDENCE

Correspondence and material requests should be addressed to David Gómez-Varela.

## ACKNOWLEDGMENTS

We thank Elisabeth Clifford (Division of Pharmacology & Toxicology, University of Vienna, Austria) for assistance during sample preparation. We thank Astrid Horn Ph.D. (Centre for Microbiology and Environmental Systems Science, Division of Microbial Ecology, Austria) for suggestions regarding bacterial counting. We Thank Vadim Demichev for the discussions related to DIA-NN analysis. We thank Biognosys AG for the 1-month free license of Spectronaut. This work is funded by a Research grant from the FWF (P35856-B to M.S.) and the University of Vienna. The computational results presented have been achieved in part using the Vienna Scientific Cluster (VSC).

---

**Supplementary Figure 1:**
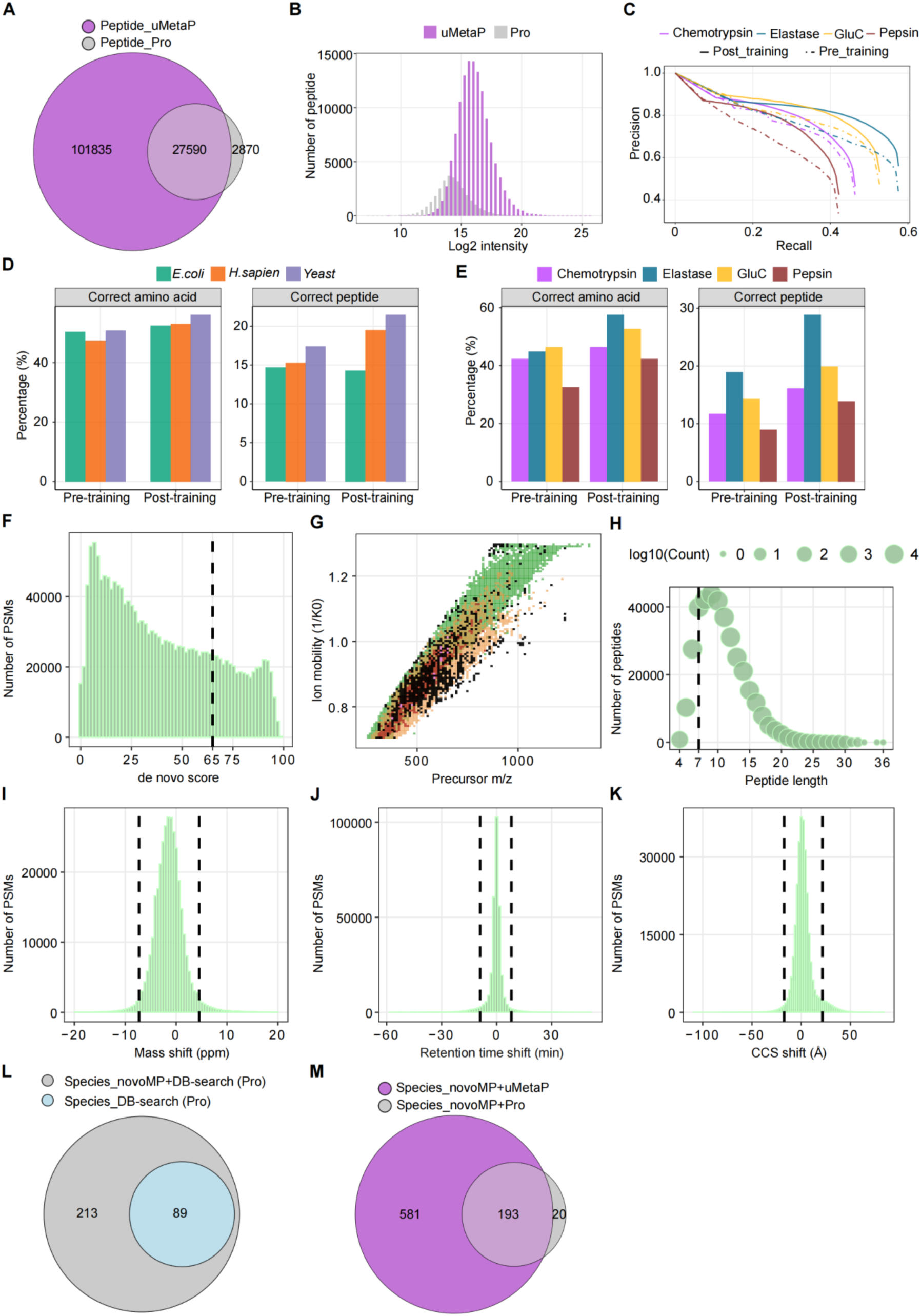
Comprehensive evaluation of novoMP performance in peptide identification and taxonomic annotation. (A) Overlap of peptides identified by current uMetaP (magenta) and previous timsTOF Pro workflows (gray) using same pre-fractionated samples. (B) Log2 intensity distribution of peptides identified by uMetaP and previous timsTOF Pro workflows. (C) Precision-recall curves comparing pre-training and post-training performance of Novor algorithm across enzymes (Chemotrypsin, Elastase, GluC, and Pepsin). (D) Percentage of correct amino acid and peptide identifications for in a dataset generated from species-mix samples (*E. coli*, *H. sapiens*, and yeast) in pre- and post-training conditions of Novor. (E) Percentage of correct amino acid and peptide identifications across datasets prepared with various enzymes. (F-K) Filtering and validation metrics applied to novoMP- derived PSMs: (F) Distribution of de novo scores, with the dotted line indicating the filtering threshold (score value = 65) for high-confidence matches. (G) Distribution of precursor charge states of de novo sequenced PSMs. Black-dots represent singly charged precursors that were excluded for further processing. (H) Stats of peptide length and corresponding counts. The black-dotted line indicates the cut-off of 7 amino acids. (I) Mass shift distribution of de novo PSMs. The black-dotted lines indicate the upper (+4.54 ppm) and lower (-7.28 ppm) cut-off to ensure 95% of the data under the distribution. (J) Distribution of retention time shifts between observed and predicted values. The black-dotted lines indicate the upper (+8.04 min) and lower (-8.99 min) cut-off to retain 95% of the data. (K) Distribution of cross-collision section (CCS) differences between observed CCS and predicted CCS. The black-dotted lines indicate the upper (+21.59 Å) and lower (-17.25 Å) cut-off to keep 95% of the data under the distribution. (L) Venn diagram showing species identified by applying novoMP (gray) and classic DB-search strategy (light blue) in a dataset acquired using our previous timsTOF Pro workflow. (M) Annotated species comparison by applying novoMP to current uMetaP workflow (magenta) and previous timsTOF Pro workflows (gray) using same pre-fractionated samples.

**Supplementary Figure 2:**
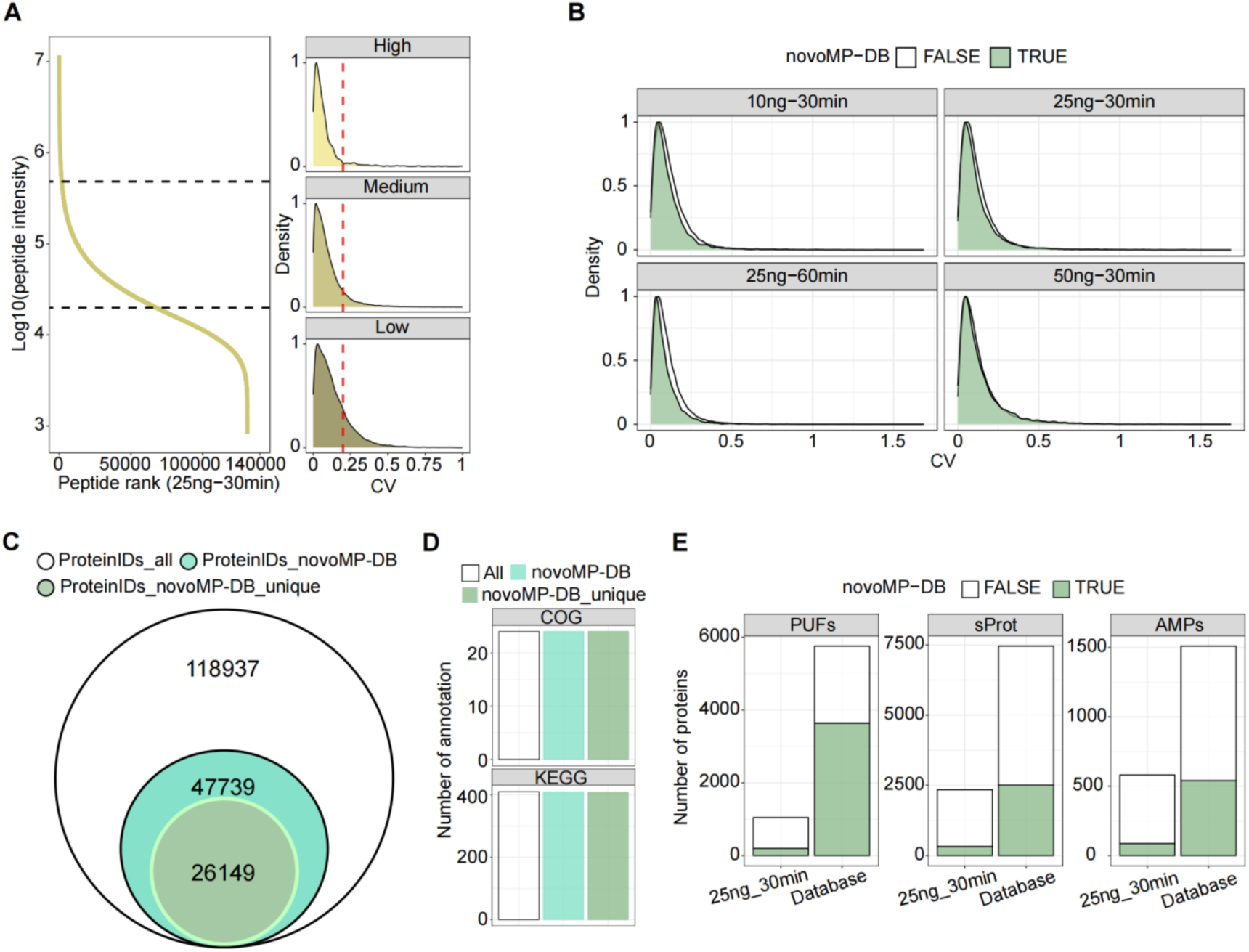
Validation of quantitative precision and functional annotations enabled by uMetaP and novoMP in complex metaproteomic datasets. (A) Peptide intensity distribution and quantitative precision analysis for 25 ng of peptides analyzed with a 30- minute LC gradient. Left: Log10 intensity distribution ranked by peptide abundance, categorized into high, medium, and low-intensity groups. Right: Density plots of the coefficient of variation (CV) for each intensity group, with the red dashed line indicating a CV threshold of 0.2. (B) Density plots of CV values (triplicates) across varying sample loadings (10 ng to 50 ng) and LC gradient lengths (30 to 60 minutes). Peptides identified by novoMP-DB (green) demonstrate comparable or superior quantitative precision to database-searched peptides across all conditions. (C) Overlap of all identified protein groups (ProteinIDs_all), those identified with novoMP-DB (ProteinIDs_novoMP-DB), and those uniquely identified by novoMP-DB (ProteinIDs_novoMP-DB_unique). (D) Functional annotations (COG and KEGG) of all identified protein groups (All), those identified with novoMP-DB (ProteinIDs_novoMP-DB), and those uniquely identified by novoMP-DB (ProteinIDs_novoMP-DB_unique). (E) Amount of PUFs, sProt, and AMPs experimentally detected using 25 ng with a 30-min gradient and present in the constructed microbial protein database.

**Supplementary Figure 3:**
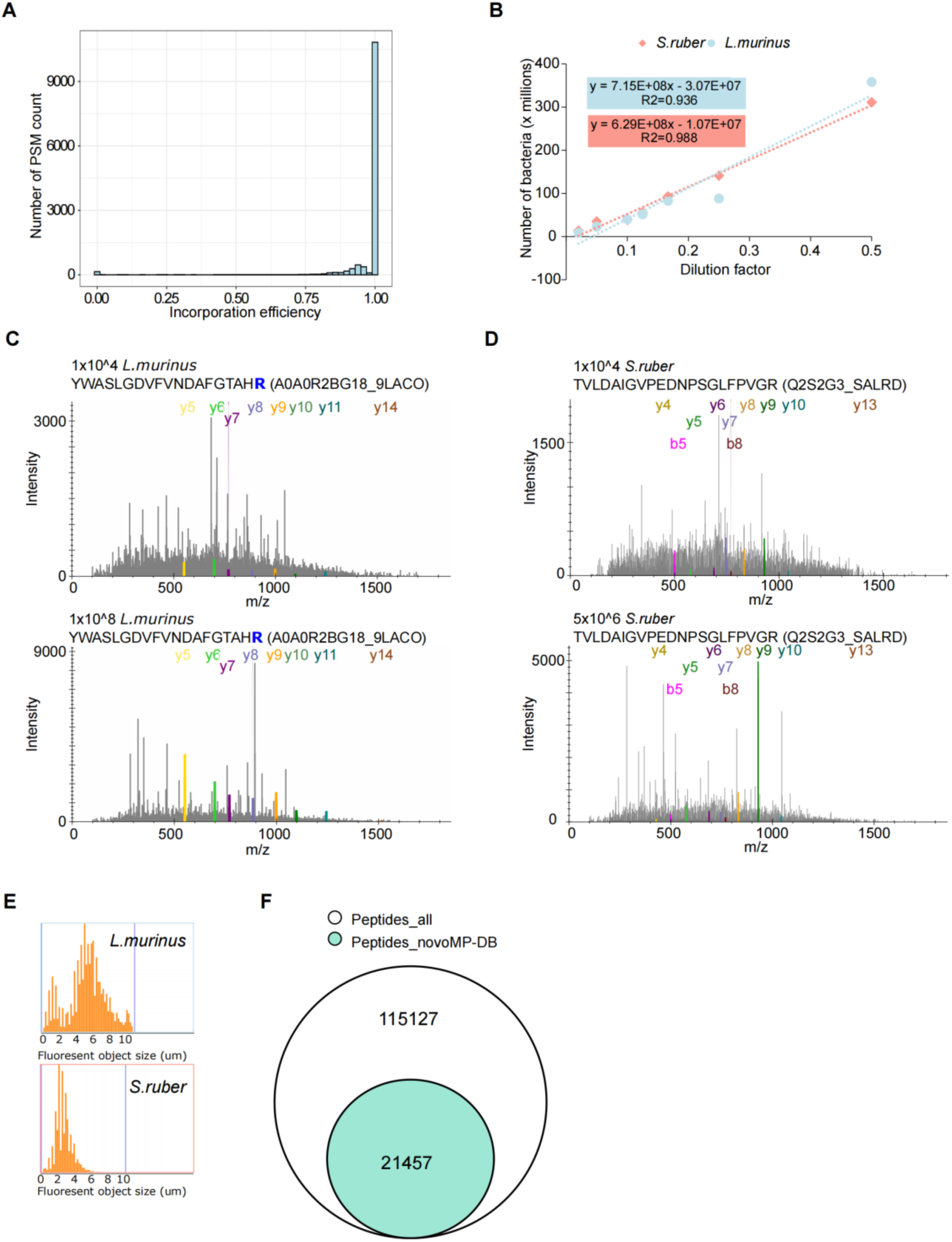
Validation and characteristics of peptides and proteins identified in spiked-in bacterial experiments. (A) Distribution of SILAC incorporation efficiency for *L. murinus*, showing an average incorporation efficiency of 97.42%. (B) Linear relationship between dilution factors and counted bactria for *S. ruber* and *L. murinus*. (C-D) Representative of identified MS/MS spectra of peptides from *L. murinus* (C) and *S. ruber* (D) at the LoD of 10^4^ bacterial cells compared to same peptides at higher spike-in amounts. (E) Fluorescent object size distribution for *L. murinus* and *S. ruber* measured during bacterial counting. (F) Venn diagram comparing the total peptides identified (Peptides_all) and those identified by novoMP-DB (Peptides_novoMP-DB).

**Supplementary Figure 4:**
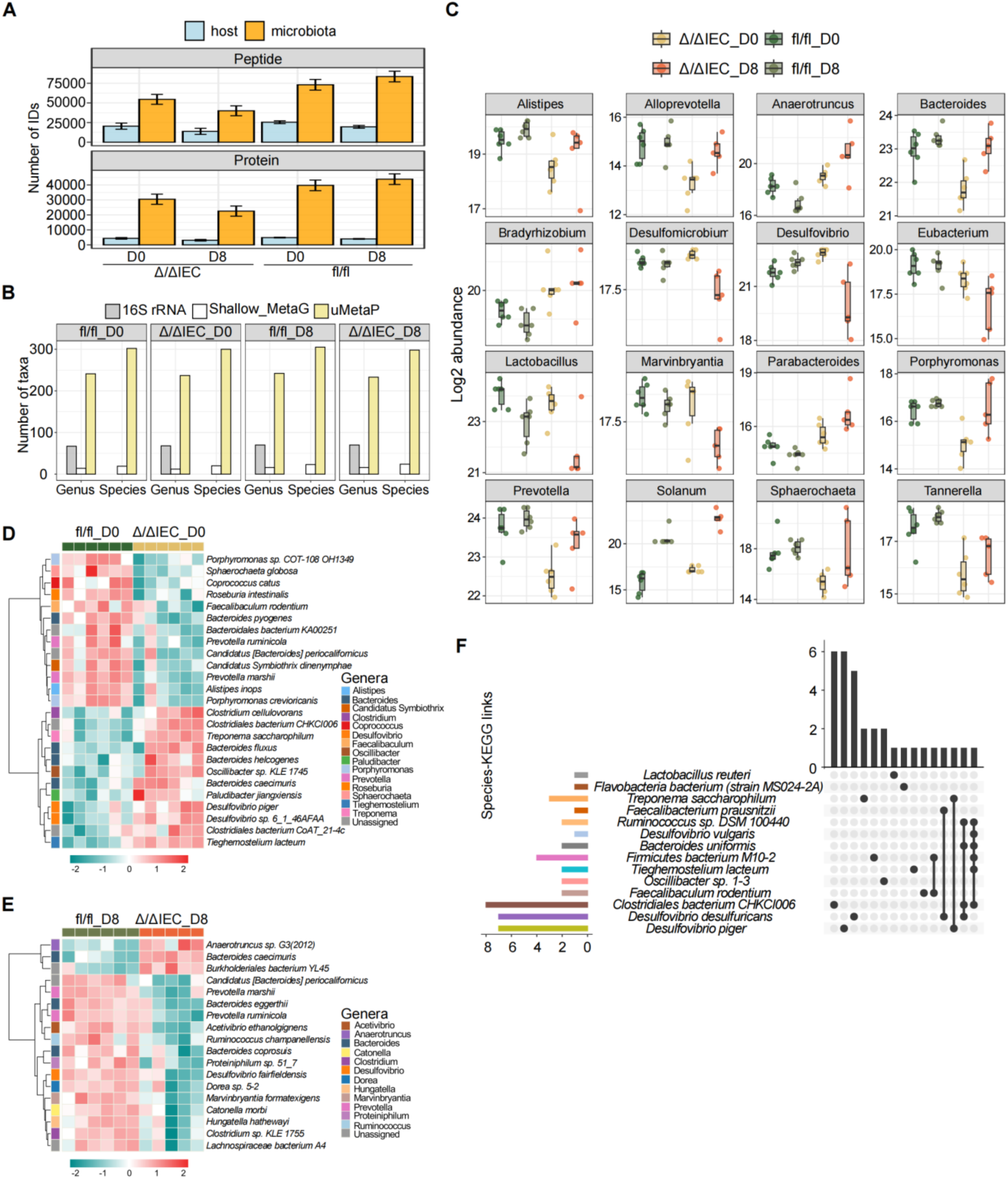
Microbial taxonomic and functional changes during intestinal injury in response to mitochondrial dysfunction. (A) Number of peptides and protein identifications for host and microbiota across D0 and D8 in control (Hsp60^fl/fl^) and injured (Hsp60^Δ/ΔIEC^) mice. (B) Comparison of detected genera and species (bacteria superkingdom) using 16S rRNA, shallow shotgun metagenomic sequencing, and uMetaP on the same sample set. (C) Log2 abundance of 16 significantly altered genera in response to metabolic injury discovered by uMetaP. (D-E) Abundances of differentially altered species at D0 (D) and D8 (E). The genera assignments of those species are colored and shown on the left of the heatmaps. (F) UpSet plot showing uniqueness and shareness of significantly regulated KEGG pathways among 14 species at D0.

**Supplementary Figure 5:**
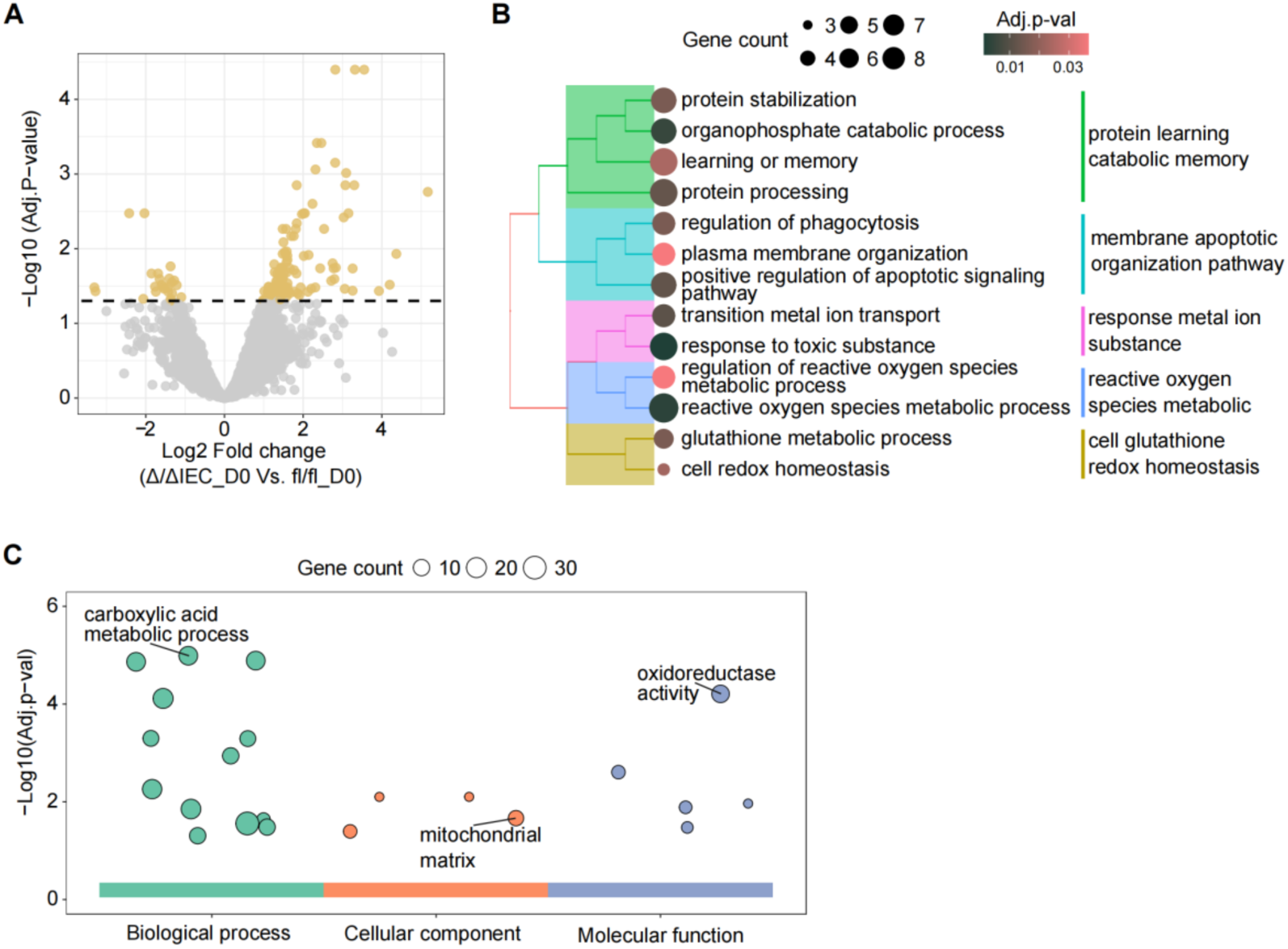
Functional enrichment analysis of host proteome changes. (A) Volcano plot showing log2 fold changes of significantly regulated proteins at D0 (Δ/ΔIEC_D0 vs. fl/fl_D0). Proteins with adjusted p-values ≤ 0.05 are highlighted. (B) Enriched biological processes from significantly regulated proteins at D0. (C) GO enrichment of 33 proteins consistently regulated in mouse metaproteomics (colonic content), mouse targeted RNA analysis (colon tissues), and human transcriptomics datasets (Crohn’s disease, ileum biopsy). The enriched terms are sized based on the number of genes mapped.

## REFERENCES

1. Clemente JC, Ursell LK, Parfrey LW, Knight R. The impact of the gut microbiota on human health: an integrative view. Cell 148, 1258–1270 (2012).

2. Martinez I, Muller CE, Walter J. Long-term temporal analysis of the human fecal microbiota revealed a stable core of dominant bacterial species. PLoS One 8, e69621 (2013).

3. Turnbaugh PJ, et al. A core gut microbiome in obese and lean twins. Nature 457, 480–484 (2009).

4. Duan H, et al. Assessing the Dark Field of Metaproteome. Anal Chem 94, 15648–15654 (2022).

5. Zhou X, et al. Longitudinal profiling of the microbiome at four body sites reveals core stability and individualized dynamics during health and disease. Cell Host & Microbe 32, 506–526.e509 (2024).

6. Van Den Bossche T, et al. Critical Assessment of MetaProteome Investigation (CAMPI): a multi-laboratory comparison of established workflows. Nat Commun 12, 7305 (2021).

7. Chapman JD, Edgar JS, Goodlett DR, Goo YA. Use of captive spray ionization to increase throughput of the data-independent acquisition technique PAcIFIC. Rapid Commun Mass Spectrom 30, 1101–1107 (2016).

8. Brunner AD, et al. Ultra-high sensitivity mass spectrometry quantifies single-cell proteome changes upon perturbation. Mol Syst Biol 18, e10798 (2022).

9. Ctortecka C, et al. Automated single-cell proteomics providing sufficient proteome depth to study complex biology beyond cell type classifications. bioRxiv, (2024).

10. Le Bihan T, et al. De novo protein sequencing of antibodies for identification of neutralizing antibodies in human plasma post SARS-CoV-2 vaccination. Nature Communications 15, 8790 (2024).

11. Gomez-Varela D, Xian F, Grundtner S, Sondermann JR, Carta G, Schmidt M. Increasing taxonomic and functional characterization of host-microbiome interactions by DIA-PASEF metaproteomics. Front Microbiol 14, 1258703 (2023).

12. Paik YK, et al. Launching the C-HPP neXt-CP50 Pilot Project for Functional Characterization of Identified Proteins with No Known Function. J Proteome Res 17, 4042–4050 (2018).

13. Petruschke H, Anders J, Stadler PF, Jehmlich N, von Bergen M. Enrichment and identification of small proteins in a simplified human gut microbiome. J Proteomics 213, 103604 (2020).

14. Sberro H, et al. Large-Scale Analyses of Human Microbiomes Reveal Thousands of Small, Novel Genes. Cell 178, 1245–1259 e1214 (2019).

15. Ma Y, et al. Identification of antimicrobial peptides from the human gut microbiome using deep learning. Nat Biotechnol 40, 921–931 (2022).

16. Tim Van Den Bossche DB, Sam van Puyenbroeck, Tomi Suomi, Tanja Holstein, Lennart Martens, Laura L. Elo, Thilo Muth. Metaproteomics beyond databases: addressing the challenges and potentials of de novo sequencing. ChemRxiv, (2024).

17. Armengaud J. Metaproteomics to understand how microbiota function: The crystal ball predicts a promising future. Environmental Microbiology 25, 115–125 (2023).

18. Ma B. Novor: real-time peptide de novo sequencing software. J Am Soc Mass Spectrom 26, 1885–1894 (2015).

19. Qin J, et al. A human gut microbial gene catalogue established by metagenomic sequencing. Nature 464, 59–65 (2010).

20. Yang J, et al. Species-Level Analysis of Human Gut Microbiota With Metataxonomics. Front Microbiol 11, 2029 (2020).

21. Taibi A, et al. Data on cecal and fecal microbiota and predicted metagenomes profiles of female mice receiving whole flaxseed or its oil and secoisolariciresinol diglucoside components. Data Brief 38, 107409 (2021).

22. Lloyd-Price J, et al. Multi-omics of the gut microbial ecosystem in inflammatory bowel diseases. Nature 569, 655–662 (2019).

23. Metwaly A, Reitmeier S, Haller D. Microbiome risk profiles as biomarkers for inflammatory and metabolic disorders. Nat Rev Gastroenterol Hepatol 19, 383–397 (2022).

24. Urbauer E, et al. Mitochondrial perturbation in the intestine causes microbiota-dependent injury and gene signatures discriminative of inflammatory disease. Cell Host & Microbe 32, 1347–1364.e1310 (2024).

25. Khaloian S, et al. Mitochondrial impairment drives intestinal stem cell transition into dysfunctional Paneth cells predicting Crohn’s disease recurrence. Gut 69, 1939 (2020).

26. Radoux CJ, Vianello F, McGreig J, Desai N, Bradley AR. The druggable genome: Twenty years later. Frontiers in Bioinformatics 2, (2022).

27. Ngollo M, et al. Identification of Gene Expression Profiles Associated with an Increased Risk of Post-Operative Recurrence in Crohn’s Disease. Journal of Crohn’s and Colitis 16, 1269–1280 (2022).

28. Chawla M, et al. An epithelial Nfkb2 pathway exacerbates intestinal inflammation by supplementing latent RelA dimers to the canonical NF-κB module. Proc Natl Acad Sci U S A 118, (2021).

29. Dhillon SS, et al. Higher activity of the inducible nitric oxide synthase contributes to very early onset inflammatory bowel disease. Clin Transl Gastroenterol 5, e46 (2014).

30. Park SC, Jeen YT. Anti-integrin therapy for inflammatory bowel disease. World J Gastroenterol 24, 1868–1880 (2018).

31. Dumas T, et al. The astounding exhaustiveness and speed of the Astral mass analyzer for highly complex samples is a quantum leap in the functional analysis of microbiomes. Microbiome 12, 46 (2024).

32. Wang A, et al. Assessing fecal metaproteomics workflow and small protein recovery using DDA and DIA PASEF mass spectrometry. Microbiome Research Reports 3, 39 (2024).

33. Sun Y, et al. metaExpertPro: A Computational Workflow for Metaproteomics Spectral Library Construction and Data-Independent Acquisition Mass Spectrometry Data Analysis. Molecular & Cellular Proteomics 23, 100840 (2024).

34. Creskey M, et al. Metaproteomics reveals age-specific alterations of gut microbiome in hamsters with SARS-CoV-2 infection. bioRxiv, 2024.2011.2012.623292 (2024).

35. Lohmann P, et al. Function is what counts: how microbial community complexity affects species, proteome and pathway coverage in metaproteomics. Expert Rev Proteomics 17, 163–173 (2020).

36. Sabine M-S, Pratik DJ, Timothy JG, Mélanie B, Ruddy W. Chapter 17 - Comparative Metaproteomics to Study Environmental Changes. In: Metagenomics (ed Muniyandi N). Academic Press (2018).

37. Wexler AG, Goodman AL. An insider’s perspective: Bacteroides as a window into the microbiome. Nature Microbiology 2, 17026 (2017).

38. Cömert C, Fernandez-Guerra P, Bross P. A Cell Model for HSP60 Deficiencies: Modeling Different Levels of Chaperonopathies Leading to Oxidative Stress and Mitochondrial Dysfunction. In: Protein Misfolding Diseases: Methods and Protocols (ed Gomes CM). Springer New York (2019).

39. Sensi P. History of the development of rifampin. Rev Infect Dis 5 **Suppl 3**, S402–406 (1983).

40. Wexler HM. Bacteroides: the good, the bad, and the nitty-gritty. Clin Microbiol Rev 20, 593–621 (2007).

41. Berger E, et al. Mitochondrial function controls intestinal epithelial stemness and proliferation. Nature Communications 7, 13171 (2016).

42. Kong AT, Leprevost FV, Avtonomov DM, Mellacheruvu D, Nesvizhskii AI. MSFragger: ultrafast and comprehensive peptide identification in mass spectrometry-based proteomics. Nat Methods 14, 513–520 (2017).

43. Kall L, Canterbury JD, Weston J, Noble WS, MacCoss MJ. Semi-supervised learning for peptide identification from shotgun proteomics datasets. Nat Methods 4, 923–925 (2007).

44. Xu T, et al. ProLuCID: An improved SEQUEST-like algorithm with enhanced sensitivity and specificity. J Proteomics 129, 16–24 (2015).

45. Tabb DL, McDonald WH, Yates JR, 3rd. DTASelect and Contrast: tools for assembling and comparing protein identifications from shotgun proteomics. J Proteome Res 1, 21–26 (2002).

46. Prianichnikov N, et al. MaxQuant Software for Ion Mobility Enhanced Shotgun Proteomics. Mol Cell Proteomics 19, 1058–1069 (2020).

47. Bouwmeester R, Gabriels R, Hulstaert N, Martens L, Degroeve S. DeepLC can predict retention times for peptides that carry as-yet unseen modifications. Nature Methods 18, 1363–1369 (2021).

48. Declercq A, et al. TIMS^2^Rescore: A DDA-PASEF optimized data-driven rescoring pipeline based on MS^2^Rescore. *bioRxiv*, 2024.2005.2029.596400 (2024).

49. Boratyn GM, et al. BLAST: a more efficient report with usability improvements. Nucleic Acids Research 41, W29–W33 (2013).

50. Buchfink B, Reuter K, Drost H-G. Sensitive protein alignments at tree-of-life scale using DIAMOND. Nature Methods 18, 366–368 (2021).

51. Demichev V, Messner CB, Vernardis SI, Lilley KS, Ralser M. DIA-NN: neural networks and interference correction enable deep proteome coverage in high throughput. Nat Methods 17, 41–44 (2020).

52. Kistner F, Grossmann JL, Sinn LR, Demichev V. QuantUMS: uncertainty minimisation enables confident quantification in proteomics. *bioRxiv*, 2023.2006.2020.545604 (2023).

53. Cox J, Hein MY, Luber CA, Paron I, Nagaraj N, Mann M. Accurate proteome-wide label-free quantification by delayed normalization and maximal peptide ratio extraction, termed MaxLFQ. Mol Cell Proteomics 13, 2513–2526 (2014).

54. MacLean B, et al. Skyline: an open source document editor for creating and analyzing targeted proteomics experiments. Bioinformatics 26, 966-968 (2010).

55. Cheng K, et al. MetaLab 2.0 Enables Accurate Post-Translational Modifications Profiling in Metaproteomics. J Am Soc Mass Spectrom 31, 1473–1482 (2020).

56. Cantalapiedra CP, Hernandez-Plaza A, Letunic I, Bork P, Huerta-Cepas J. eggNOG-mapper v2: Functional Annotation, Orthology Assignments, and Domain Prediction at the Metagenomic Scale. Mol Biol Evol 38, 5825–5829 (2021).

57. Porcheddu M, Abbondio M, De Diego L, Uzzau S, Tanca A. Meta4P: A User-Friendly Tool to Parse Label-Free Quantitative Metaproteomic Data and Taxonomic/Functional Annotations. Journal of Proteome Research 22, 2109–2113 (2023).

58. Wu T, et al. clusterProfiler 4.0: A universal enrichment tool for interpreting omics data. *The Innovation* 2, (2021).

59. Cannon M, et al. DGIdb 5.0: rebuilding the drug–gene interaction database for precision medicine and drug discovery platforms. Nucleic Acids Research 52, D1227–D1235 (2024).

